# The mevalonate pathway of isoprenoid biosynthesis supports metabolic flexibility in *Mycobacterium marinum*

**DOI:** 10.1101/2025.07.11.664281

**Authors:** Christine M. Qabar, Edward E.K. Baidoo, Emine Akyuz Turumtay, Tariq M. Qayum, Jay D. Keasling, Cressida A. Madigan, Daniel A. Portnoy, Jeffery S. Cox

## Abstract

Isoprenoids are a diverse class of natural products that are essential in all domains of life. Most bacteria synthesize isoprenoids through either the methylerythritol phosphate (MEP) pathway or the mevalonate (MEV) pathway, while a small subset encodes both pathways, including the pathogen *Mycobacterium marinum* (Mm). It is unclear whether the MEV pathway is functional in Mm, or why Mm encodes seemingly redundant metabolic pathways. Here we show that the MEP pathway is essential in Mm while the MEV pathway is dispensable in culture, with the ΔMEV mutant having no growth defect in axenic culture but a competitive growth defect compared to WT Mm. We found that the MEV pathway does not play a role in *ex vivo* or *in vivo* infection but does play a role in survival of peroxide stress. Metabolite profiling revealed that modulation of the MEV pathway causes compensatory changes in the concentration of MEP intermediates DOXP and CDP-ME, suggesting that the MEV pathway is functional and that the pathways interact at the metabolic level. Finally, the MEV pathway is upregulated early in the shift down to hypoxia, suggesting that it may provide metabolic flexibility to this bacterium. Interestingly, we found that our complemented strains, which vary in copy number of the polyprenyl synthetase *idsB2*, responded differently to peroxide and UV stresses, suggesting a role for this gene as a determinant of downstream prenyl phosphate metabolism. Together, these findings suggest that MEV may serve as an anaplerotic pathway to make isoprenoids under stress conditions.

**Importance:** Organisms from all domains of life utilize isoprenoids to carry out thousands of critical and auxiliary cellular processes, including signaling, membrane integrity, stress response, and host-pathogen interactions. The common precursor of all isoprenoids is synthesized via one of two biosynthetic pathways and importantly, some bacteria encode both pathways, including *M. marinum*. We found that only one pathway is essential in *M. marinum*, while the nonessential pathway may confer metabolic flexibility to help the bacterium better adapt to various environmental conditions. We also found that the polyprenyl synthetase IdsB2 plays an important role in driving such phenotypes. Further, we demonstrate metabolic interplay between both functional pathways. These insights represent the first characterization of isoprenoid biosynthesis in dual pathway-encoding mycobacteria.

## Introduction

Isoprenoids are the largest class of natural products, of which 95,000 have been identified to date (1, 2). In bacteria, isoprenoids play diverse and critical roles, spanning microbial signaling, stress response, electron transport, and construction of the cell wall (3). Despite wide structural and functional diversity, all isoprenoids arise from the fundamental precursor isopentenyl pyrophosphate (IPP) and its isomer dimethylallyl pyrophosphate (DMAPP). These terpenoid building blocks can be synthesized via two independent pathways: the mevalonate (MEV) pathway, which is primarily found in eukaryotes and archaea, and the methylerythritol phosphate (MEP) pathway, which is found in bacteria, plastids, and algae (4). Downstream of IPP are a series of condensation reactions shared by both the MEV and MEP pathways, collectively known as prenyl phosphate metabolism.

While the MEV and MEP pathways both produce IPP, they consume different starting reagents and proceed via unique enzymatic steps which produce distinct intermediates (Figure 1A). All eukaryotes and archaea use the MEV pathway for isoprenoid biosynthesis, but there is much diversity in pathway utilization among bacteria. While most bacteria exclusively use the MEP pathway, some bacteria use only the MEV pathway, and others yet encode both pathways (5). Some obligate intracellular bacteria, like *Rickettsia parkeri*, lack any endogenous pathway and instead rely on import of host IPP for metabolism (6).

**Figure 1:**
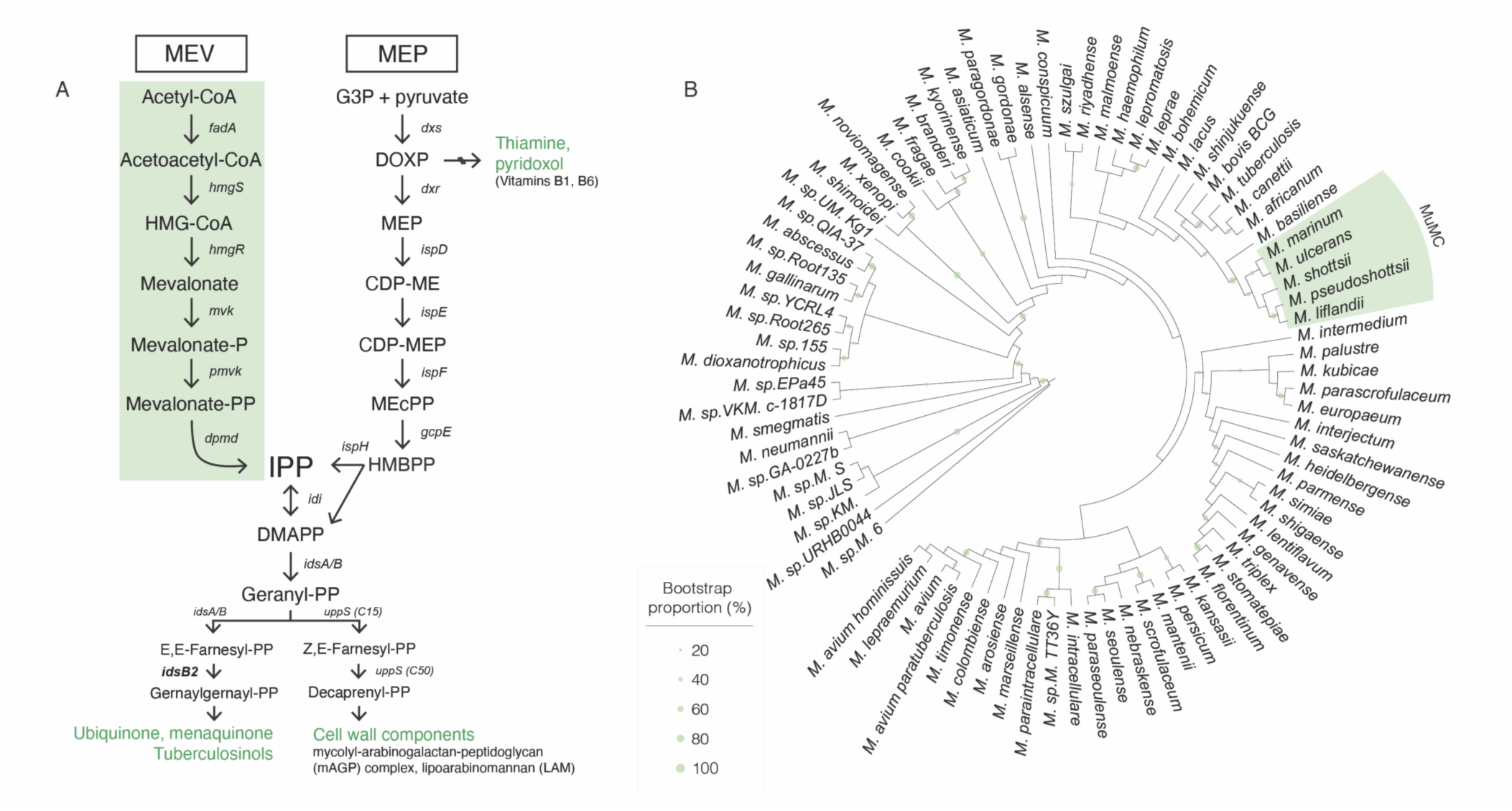
Mm encodes two pathways for isoprenoid biosynthesis. **A. Two pathways of isoprenoid biosynthesis.** Diagram of the MEV and MEP pathways of isoprenoid precursor biosynthesis. Enzymatic steps are indicated by italicized gene names. Important end products in mycobacteria are shown in green. **B. MEV distribution in mycobacteria.** The *M. ulcerans-M. marinum* (MuMC) clade is the only group of mycobacteria that encodes the MEV pathway. All mycobacteria, including the MuMC clade, encode the MEP pathway. Shading in green indicates species that encode MEV genes. Tree represents 84 mycobacterial species; sequences and alignment data are available in Table S1 and File S1.

Insights into the origins of these different pathways can be gleaned from comparative genomics. MEP genes are scattered throughout the genomes of MEP-encoding bacteria and conversely, MEV genes are found within a single operon, leading some to speculate acquisition via horizontal gene transfer (7, 8). Phylogenomic analyses suggest that the punctate distribution of MEV genes among bacteria can be traced back to a cenancestral MEV pathway, and subsequently some bacterial lineages lost these genes as they evolved the MEP pathway (9). Mycobacteria present a unique divergence from this hypothesis, given that only one clade of mycobacteria encodes both pathways, including the marine pathogen *Mycobacterium marinum* (Mm). The evolutionary pressure promoting retention of the MEV pathway among certain bacteria remains unknown, however horizontal gene transfer is not likely for Mm, as the GC content of the MEV operon is 64.56% compared to 65.73% for the rest of the genome (10), suggesting that this pathway was not recently acquired.

In bacteria encoding both MEP and MEV genes, there is evidence that these pathways are nonredundant and may support metabolic flexibility to a variety of stressors. Members of the *Actinomycetota* phylum, which also contains the genus *Mycobacterium*, use the MEP pathway early in growth for primary metabolism and later activate the MEV pathway to synthesize secondary metabolites (8). Interestingly, intermediates in the pathways have been implicated as effectors themselves, including the MEP metabolite (E)-4-hydroxy-3-methyl-but-2-enyl pyrophosphate (HMBPP), which is the most potent activator of host Vγ9Vδ2 T cells (11–13). Another compelling line of evidence points to the MEP pathway as an oxidative stress sensor, as the final two enzymes in this pathway contain oxygen-reactive iron-sulfur clusters (14, 15). Further, the MEP pathway intermediate 2-C-methyl-D-erythritol-2,4-cyclopyrophosphate (MEcPP) accumulates in response to oxidative stress in bacteria and is a stress signaling molecule in plastidal plants (16, 17), supporting a model in which encoding both pathways may confer resistance to environmental stresses.

One interesting case is in *Listeria monocytogenes* (Lm), a Gram-positive intracellular pathogen, which also encodes both pathways. Lm relies on the MEV pathway for metabolism, yet encodes an MEP pathway that is aerobically nonfunctional due to the failure of the final two genes in the pathway (18). The related *Listeria innocua* encodes an incomplete MEP pathway, including all but those same two final MEP genes which are directly downstream of the synthesis of MEcPP (19). Further, a mutant in the MEcPP synthase *ispF* was identified as more sensitive to oxidative stress in Lm (20). Taken together, it is highly likely that the MEP pathway in *Listeria* synthesizes the putative oxygen sensor MEcPP, while the MEV pathway primarily supports isoprenoid metabolism. However this remains to be explored in Mm, which also encodes both pathways.

The MEP pathway is essential in the important human pathogen *M. tuberculosis*, its vaccine strain *M. bovis* BCG, and likely all mycobacteria that exclusively use the MEP pathway; however, it is unclear if one or both pathways is essential in Mm (21, 22). Previous transposon mutagenesis experiments failed to identify several genes in the MEP pathway as essential in Mm (23, 24). In contrast, here we provide conclusive genetic evidence of the essentiality of the MEP pathway in Mm and, conversely, the dispensability of the MEV pathway. Further, in this work we demonstrate that although the MEV pathway is nonessential, it is intact, functional, and supports the ability of Mm to persist in dynamic environmental conditions.

## Methods

### Bioinformatic analysis

Phylogenetic tree building was done as previously described (25). Briefly, full-length 16s rRNA sequences were collected from one representative genome for each unique species of mycobacteria available on BioCyc (26) and the NCBI Gene database (27), representing 84 mycobacterial species. Geneious Prime (v2024.0.7) was used to perform a MUSCLE alignment (28) and build a PhyML tree (29) with 100 bootstraps. Annotation and tree visualization were performed on iTol (30). Sequences and alignment data are available in Table S1 and File S1. *In silico* SigF binding motif analysis was performed on the region upstream of the MEV operon for the consensus SigF binding motif GTTC[N17]GGGTAT between −10 and −200 of the coding region, allowing for 2 mismatches, and found one match: GTTC[agtagcggttgacgatt]TGGTAG −83bp from the start codon of *idi*. Structural and sequence alignments for IdsB2 homologs in Mm and *M. tuberculosis* were performed using MUSCLE and visualized using FastTree 2.1.11 (31). AlphaFold 3 (32) was used to predict structures, and structural alignment was performed using PyMOL (33).

### Bacterial strains and culture

*Mycobacterium marinum* M (ATCC BAA-535) was routinely grown at 30°C in Middlebrook 7H9 liquid medium or 7H10 agar (BD Difco) supplemented with 10% OADC (oleic acid-albumin-dextrose-catalase) and 0.2% Tween80. *E. coli* strain DH5α was grown at 37°C in Luria broth or agar and used for plasmid cloning. When required, the following antibiotics were used: kanamycin (25 µg/ml for Mm, 50 µg/ml for *E. coli*); hygromycin (50 µg/ml for Mm, 150 µg/ml for *E. coli*); zeocin (100 µg/ml for Mm, 50 µg/ml for *E. coli*); anhydrotetracycline (200 ng/ml for CRISPR, 500 ng/ml for ORBIT).

### Molecular cloning

Primers, oligos, and plasmids used in this study are listed in Supplementary Tables 2 and 3. Standard electroporation protocols were used for the transformation of plasmids into *E. coli*. Electrocompetent mycobacteria were prepared by washing OD_600_ 0.5-0.8 cultures four times in decreasing volumes of sterile 10% glycerol, for a final resuspension concentrated ∼20-25x. 200 µl of electrocompetent cells were combined with 5µl of plasmid DNA in a 0.2-cm electroporation cuvette and electroporated with a single pulse at 2.5 kV with 25 μF capacitance and 1,000 Ω resistance. Transformants were recovered overnight at 30°C in 7H9 medium and plated on selective 7H10 agar containing antibiotics. 3-6 clones were randomly selected and verified via PCR.

### ORBIT

Gene replacement was performed via the <u>o</u>ligonucleotide-mediated recombineering followed by <u>B</u>xb1 integrase targeting (ORBIT) system as previously described (34). Targeting oligos were designed such that an *attP* site was flanked on either side by 30 bp arms homologous to the flanking regions of the genomic region of interest. Strains were transformed with the integrase/annealase plasmid pKM444, induced with anhydrotetracycline (ATc), and transformed with both the targeting oligo and “payload plasmid” pKM464 or pKM496. Transformant colonies were screened via colony PCR and sequencing of the insertion junctions. Integrated payload plasmids were cured out of the genome with the excisionase plasmid pKM512. Transformants were induced with ATc, and the plasmid was counter selected for on 7H10 + 10% sucrose.

### CRISPRi and CRISPRn

Guide RNA sequences were designed in CHOPCHOP (35) using previously characterized PAM sequences (36). Guides were designed to be 21-24nt with the most PAM-distal nucleotide an A or G. BsmBI overhangs were added to each spacer to mediate insertion into pLJR965. Gene silencing was performed using the site-specific transcriptional repression system CRISPRi as previously described (36). Briefly, guide oligos were annealed and ligated into the integrating, dCas9-containing pLJR965 vector and the subsequent plasmid was transformed into mycobacteria. Inducible transcriptional repression was achieved by treatment with 200 ng/ml ATc. To test strains, mid-log cells were OD_600_ matched to 0.3, serially diluted in 7H9, and 3 µL spots were plated on both 7H10 agar supplemented with kanamycin alone (Kan25 µg/mL; no induction) or kanamycin and ATc (ATc200 ng/mL; induced). Gene deletion was performed using the site-specific gene disruption system CRISPRn as previously described (37). Briefly, up to two guide oligos were cloned into the Cas9-containing vector pCRISPRx, and the subsequent plasmid was transformed into mycobacteria and cells were allowed to recover overnight in 7H9. Transformants were then induced with 75 ng/ml ATc for one hour and plated on selective media. Subsequent transformants were screened via sequencing to identify successful mutants. Indels were most common, but deletions were prioritized.

### RNA isolation + RT-PCR

25 ml of OD_600_ 0.3-0.4 cultures were resuspended in 1 ml TRIzol™ (Invitrogen, catalog #15596026) and lysed via bead-beating thrice in 30 second increments, incubating on ice in between bursts. RNA was then extracted from the supernatant via chloroform-ethanol extraction, washed using PureLink™ RNA Mini wash buffers I & II (Invitrogen, catalog #12183018A), and treated with DNAse I (NEB). Following a 10 minute DNAse inactivation at 75°C, RNA was stored at −80°C. Primers were designed with the following criteria: 17-25 bp in length, 3’ G/C clamp, 75-150 bp amplicon, Tm ∼60°C, and analyzed for secondary structure. cDNA was prepared using a SuperScript™ III First-Strand Synthesis Kit (Invitrogen, catalog #18080051) and SsoAdvanced Universal SYBR Green Supermix (BioRad, catalog #1725271) was used for RT-PCR reactions on genes of interest as well as 16S for normalization. 1:10-diluted cDNA samples were run in technical triplicate on a CFX Connect Real-Time System (BioRad) running CFX Manager software. Normalized expression ratios were obtained via the 2^-ΔΔCq^ (Livak) Method (38).

### Plasmid loss frequency analysis

Strains were passaged every 48 hours in fresh 7H9 media without antibiotic selection, serially diluted, and plated on both plain and hygromycin-containing 7H10 agar. After 7 days incubation at 30°C, dilutions were counted, and CFU/ml was calculated for each dilution. Percent loss is calculated as [(CFU/ml on plain 7H10)-(CFU/ml on hyg50)]/(CFU/ml on plain 7H10).

### Growth curve

Cultures were grown to mid-log (OD_600_ 0.5-0.8) and back diluted to 0.05 in 7H9. 100 μl of each strain was loaded to respective wells of 96-well flat-bottom microplate (ThermoFisher, catalog #267427). To prevent evaporation of the sample wells, all remaining wells were filled with uninoculated 7H9 and the exterior troughs were filled with 1% agarose. Cultures were shaken at 30°C and optical density (OD_600_) readings were taken every 2 hours for 8 days on a BioTek Epoch 2 Microplate Spectrophotometer (Agilent) running Gen5 software (v3.11). Readings were normalized to blank 7H9 controls and OD-corrected. Data points acquired during logarithmic growth (determined to be between 38-54hr) were fit with a linear regression, giving the equation y=*k*X+logY_0_. The rate constant *k* was used to determine the doubling time using the equation t = ln^2/k^.

### Metabolite extraction + metabolomic analysis

The metabolite extraction protocol was modified from previously described mycobacterial extraction (39). Briefly, cells were grown to a target OD_600_ of 0.2 (early log), 0.5 (mid log), 1 (late log), or 2 (early stationary) and the equivalent of 10 OD_600_ units were pelleted, resuspended in metabolite extraction buffer (2:2:1 methanol:acetonitrile:water), and lysed via bead beating six times in 30 second increments at 4°C. Lysate was then pelleted, and supernatant was fractionated through 3 kDa ultra centrifugal filter columns (Amicon, catalog #UFC500324). Extracts were held at −80°C until targeted metabolomic analysis. Liquid chromatography–mass spectrometry (LC-MS) analysis of metabolites was performed as previously described (40). Adenylate energy charge (AEC) was calculated using (ATP + ½ ADP)/(AMP + ADP + ATP) (41). OD_600_ 0.2, 0.5, and 2 samples were isolated and analyzed in a different experiment from OD_600_ 1 samples. All metabolomics data are available in File S3.

### Competitive co-culture

Cultures were grown to mid-log (OD_600_ 0.5-0.8) and back diluted to 0.05 in 7H9 a day prior to inoculating the co-cultures to ensure uniform growth stage across strains. Strains were inoculated into co-culture tubes at an individual OD_600_ of 0.04 and either transferred to plastic inkwells (normoxic growth) or sealed in tubes with rubber stoppers per the Wayne model (hypoxic growth). Cultures were shaken at 120 rpm (normoxic) or stirred at 360 rpm (hypoxic) at 30°C for the entire time course. Timepoints were taken daily from day 0 to day 7 (normoxic) or every other day from day 0 to day 12 (hypoxic) and plated on both kanamycin- or hygromycin-containing plates. CFU from each plate were counted on day 5 after plating. Normalized competitive index was calculated using [(mutant CFU/ml)/(WT CFU/ml)]*(input CI).

### Growth *in spent media*

Cultures of either strain individually (axenic) or both strains together (co-culture) were grown to OD_600_ 1.6 (axenic) or 2.8 (co-culture), pelleted, and the supernatant was sterilized through a 0.22 µm filter, resulting in “spent media”. Prior to inoculating new cultures, spent media was supplemented with 0.002 g/ml dextrose and 0.5% glycerol to replenish critical nutrients. Supplemented spent media was then inoculated with WT Mm and OD_600_ and CFU/ml were measured as described above.

### Ex vivo macrophage infection

Murine bone marrow-derived macrophages (BMMs) were cultured in DMEM (Gibco) supplemented with 10% FBS, 10% M-CSF supernatant produced by 3T3-MCSF cells as previously described, 2 mM L-glutamine (Gibco), and 1 mM sodium pyruvate (Gibco). BMMs were plated at a density of 3×10^4^ cells per well of a 96-well plate and incubated at 37°C for 24hr. Cells were then treated with 2ng/µl IFNγ and incubated at 37°C for 24hr. On the day of infection, the bacterial inoculum was prepared as follows: mid-log (OD_600_ 0.6-0.8) mycobacterial cultures were washed in 1X D-PBS + 0.01% Tween80 and an MOI of 1 inoculum was prepared in BMM media supplemented with 10% horse serum for opsonization. BMMs were then spinfected by the opsonized bacteria and incubated at 37°C for 10 minutes to allow for phagocytosis. Following phagocytosis, supernatant was removed and replaced with fresh BMM media. Plates were incubated at 33°C with daily media changes. At each timepoint, plates were fixed and washed thrice in 1X D-PBS. Cells were stained with 1 µg/ml DAPI (Invitrogen, catalog #D1306) immediately before imaging on the Opera Phenix High-Content Screening System. Images were analyzed using Harmony high-content imaging and analysis software to quantify fluorescent area of bacteria (mCherry) and number of nuclei (DAPI).

### Zebrafish infection

Zebrafish husbandry and experiments were performed in compliance with the United States National Institutes of Health and approved by the University of California, San Diego Institutional Animal Care and Use Committee. Wild type AB strain zebrafish larvae were incubated at 28.5°C in fish water as previously described (42). Larvae were anesthetized with 2.8% Syncaine (Syndel, catalog #886862) prior to infection and imaging. Larvae of both sex were injected in the caudal vein at 3 days post fertilization using a glass capillary needle with bacterial suspensions diluted in phenol red to allow visualization of successful injection. To determine input CFU, the same infection inoculum was plated directly onto 7H10 agar supplemented with kanamycin (25 µg/ml). Input CFU was optimally determined to be ∼40-60 CFU per larva. To normalize bacterial burden across bacterial strains, larvae were infected with varying inocula of each strain (n ≥ 10 larvae per bacterial strain at 1:10 or 1:15 dilutions) and fluorescence pixel count (FPC) analysis (42) was used to inoculum match across bacterial strains. Optimal dilutions were determined to be 1:10 for WT Mm and 1:15 for ΔMEV. At 2 and 3 days post infection, zebrafish were transferred to an optical 96-well plate and imaged on a Nikon Eclipse Ti2 microscope. ImageJ was used to determine FPC and brain dissemination.

### Growth in self-generated hypoxia

Hypoxic growth curves were performed in accordance with the Wayne model of nonreplicating persistence as previously described (43). Briefly, mid log (OD_600_ 0.50) cultures were back diluted tenfold and transferred into glass tubes with 8mm stir bars (Grainger, catalog #401R24) to a headspace ratio of 0.5. Cultures were then sealed with a rubber stopper (ThermoFisher, catalog #FB57875) and incubated at 30°C with stirring. OD_600_ and CFU samples were taken daily for a minimum of 10 days. To avoid reaeration during the time course, one tube was designated per timepoint.

### UV tolerance assay

Cultures were grown to mid-log (OD_600_ 0.5-0.8) and serially diluted in 7H9 media. 5µL of each dilution was plated on 7H10 agar and exposed at 0, 10, and 20 mJ of UV in a Stratalinker UV 1800 Crosslinker (Stratagene). CFU were counted following 7 days of growth at 30°C.

### Oxidative stress survival assay

For H_2_O_2_ survival experiments, Mm was grown in media supplemented with 10% ADS (albumin-dextrose-saline) rather than OADC which contains catalase. Mm was treated with hydrogen peroxide (H_2_O_2_, ThermoFisher catalog #BP2633500) as previously described (44). Mid-log (OD_600_ 0.6-0.8) bacteria were washed in PBS containing 0.2% Tween80, back diluted to OD_600_ 0.1, and treated with 0, 1, or 2.5 mM H_2_O_2_ for 6hr prior to serial dilution and plating for CFU. Plates were incubated at 30°C for a minimum of 6 days before counting CFU.

## Results

### The MEP pathway is essential in Mm

While most mycobacteria encode the MEP pathway exclusively, there exists a unique subset which also encodes MEV genes (Figure 1A), including the pathogen *Mycobacterium marinum*, although it was previously unknown whether the MEV pathway is functional (10). Further, species of the *M. ulcerans-M. marinum* (MuMC) clade, a cluster of highly-related subspecies that evolved from Mm, also encodes MEV genes (Figure 1B) (45–47). However the *M. ulcerans* MEV pathway is predicted to be nonfunctional due to disruption by mobile genetic elements (48), but Mm appears to encode an intact MEV pathway. All other mycobacteria, including the important human pathogen *M. tuberculosis* and its vaccine strain BCG, only encode the MEP pathway (21, 22).

To test whether the MEP pathway is essential in Mm, we attempted to make knockout strains that lacked either of two genes that encode critical enzymes in the MEP pathway: Dxr, which catalyzes the first committed step in the pathway, and IspH, which catalyzes the final reaction in the pathway (Figure 1A). We were unable to delete either gene, suggesting that the MEP pathway is essential for growth in Mm as has been reported for other mycobacteria (21, 22, 49). Using a tetracycline-inducible CRISPR interference (CRISPRi) system to conditionally inhibit expression of these two MEP genes (36), we found that transcriptional repression of *dxr* or *ispH* inhibited growth (Figure 2A,B), indicating that the MEP pathway is likely essential in Mm.

**Figure 2:**
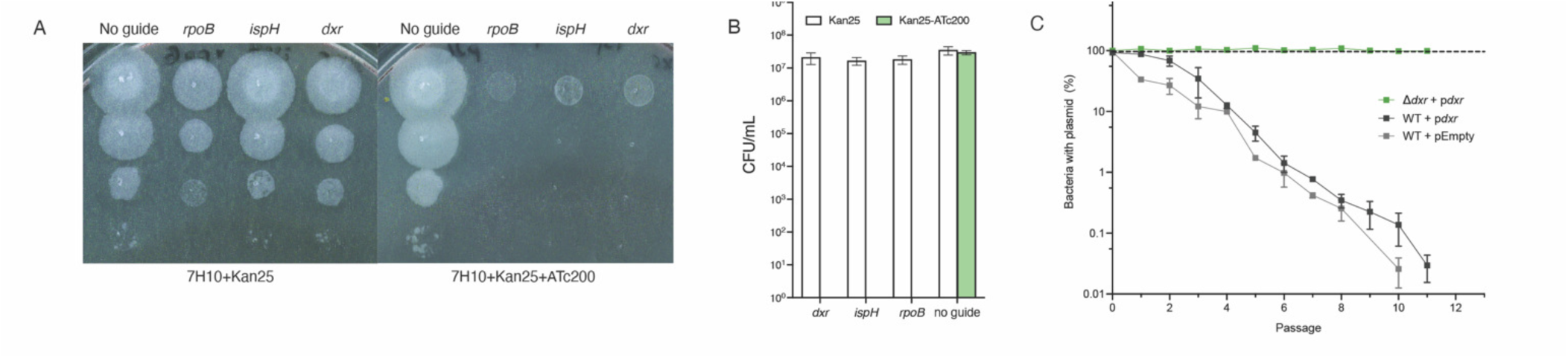
The MEP pathway is essential in Mm. **A-B. Transcriptional repression (CRISPRi) of *dxr, ispH* is lethal.** Guides targeting genes were cloned into the kanamycin-resistant vector pJLR965 as previously described (35, 36). **(A)** Upon induction of CRISPR interference machinery by addition of ATc, target genes are transcriptionally repressed. **(B)** Quantified CFU data from CRISPRi plates +/- ATc200. Data are compiled from 5 biological replicates. **C. Plasmid loss frequency analysis of a Mm mutant in Dxr, the non-mevalonate (MEP) pathway of isoprenoid biosynthesis.** MmΔ*dxr* +p*dxr*, WT Mm +p*dxr* and WT Mm +pEmpty (empty vector) were passaged every 48hr in fresh 7H9 media without antibiotics. At each passage, cultures were serially diluted and plated on both plain and hygromycin-containing agar. Loss % < 0 indicates more CFU on hyg. N=3 biological replicates.

Since we were unable to delete *dxr* in the wild-type background, we generated a merodiploid strain in which a second copy of *dxr* was expressed ectopically from an episomal plasmid (Δ*dxr* +p*dxr*). Using this recombinant strain, we were able to delete the chromosomal copy of *dxr*. To definitively test whether *dxr* is essential for growth, we performed plasmid loss experiments with the merodiploid strain. Because the plasmid vector is not faithfully transmitted to both daughter cells during every bacterial cell division (50), growth in the absence of selective pressure to maintain the plasmid will lead to progressive plasmid loss in a population. Strains were serially passaged in non-selective media over 50 generations and plated for CFU to assess plasmid loss. WT cells carrying p*dxr* were able to lose the plasmid and survive, indicating that the p*dxr* plasmid is not essential in this background (Figure 2C). In contrast, Δ*dxr* +p*dxr* had no apparent plasmid loss over the course of the experiment (Figure 2C), demonstrating that the *dxr* cargo on the plasmid is essential in the Δ*dxr* background. Together with our inability to delete *dxr* and *ispH* in wild-type Mm (data not shown), these data demonstrate that the MEP pathway is essential in Mm.

Finally, in agreement with findings in *M. tuberculosis*, we found that the Mm genome contains two non-redundant isoforms of *ispH*, also referred to in the literature as *lytB*. We were able to delete *lytB1*/MMAR_0227 (data not shown), which is homologous to the nonessential isoform in *M. tuberculosis* (51), while targeting of *ispH*/MMAR_4355, which is homologous to the essential *lytB2* in *M. tuberculosis*, was lethal (Figure 2A). Together, we demonstrate that *ispH* and *lytB1* are non-redundant in Mm.

### The MEV pathway is nonessential in Mm

Unlike the genes of the MEP pathway, the entire MEV pathway is encoded in one operon (Figure 3A). All six genes in the operon encode proteins that are clear homologs to known enzymes in the MEV pathway, and in particular the rate-limiting enzyme HmgR has sequence-level homology to known functional enzymes (Table S4, File S2). Of note, *idsB2* is the last gene in the MEV operon and catalyzes an enzymatic step downstream of IPP production that is shared by both the MEP and MEV biosynthetic pathways (Figure 1A). To test whether the MEV pathway is essential in Mm, we attempted to delete only the genes unique to the MEV pathway using <u>o</u>ligonucleotide-mediated recombineering followed by <u>B</u>xb1 integrase targeting (ORBIT; Figure 3A) (34). We successfully generated a knockout strain in which the MEV operon is deleted, then “cured” the knockout strain to avoid polar effects from the integrated ORBIT plasmid (Supplementary Figure 1A). The successful deletion of the MEV operon, hereby referred to as ΔMEV, demonstrates that the MEV pathway is not essential in Mm under standard culture conditions. We then measured the expression of *hmgR* and *dxr*, which represent the rate-limiting step of the MEV pathway and the first committed step in the MEP pathway, respectively. We observed that ΔMEV and WT had similar levels of *dxr* expression and confirmed that ΔMEV does not express *hmgR* (Figure 3B). Importantly, this also demonstrates that the MEV operon is transcriptionally active in WT Mm.

**Figure 3:**
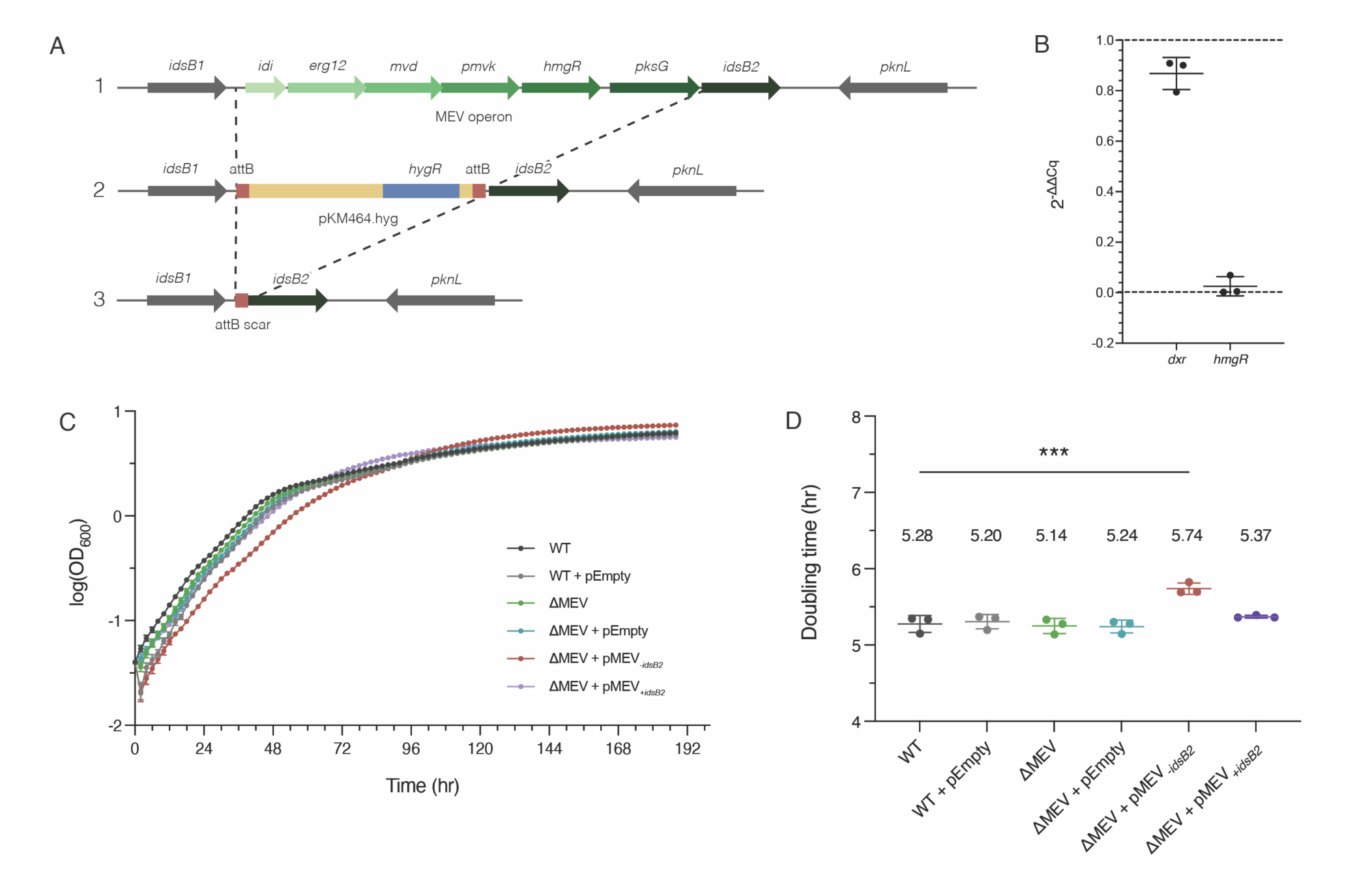
The MEV pathway is nonessential in Mm. **A. Deleting the mevalonate pathway of isoprenoid biosynthesis.** The MEV locus in Mm (step 1) was replaced with a plasmid containing a hygromycin resistance marker via homologous recombination (step 2). The gene *idsB2* was not deleted to avoid disruptions of isoprenoid synthesis downstream of IPP production. To generate a markerless deletion strain, the integrated plasmid was excised from the locus via induction of a phage excisionase enzyme (step 3). **B. Relative expression of mevalonate (MEV) and non-mevalonate (MEP) pathway genes**. Data shown are 2^-ΔΔCq^ of ΔMEV compared to WT. Sample quantification cycle (Cq) values were normalized to respective 16s controls and ΔΔCq values were calculated for representative genes in each pathway (*dxr* for MEP, *hmgR* for MEV) in the ΔMEV strain compared to WT. N = 3. **C,D. MEV deletion mutant has similar growth kinetics as WT, and complementation without *idsB2* impairs growth.** MmΔMEV has similar growth kinetics (**C**) and doubling time (**D**) to WT Mm, demonstrating that the MEV pathway does not support standard growth in culture. The complemented strain lacking *idsB2* (ΔMEV +pMEV_-*idsB2*_) has a significantly slower growth rate (**D**) compared to the other strains (***p=0.0001; one-way ANOVA and Dunnet’s multiple comparisons). pEmpty indicates empty vector control. Data plotted are mean ± SEM. N = 3 biological replicates.

To avoid disrupting global isoprenoid metabolism, we deleted the entire MEV operon except for *idsB2*, which acts downstream of both pathways. Instead, the chromosomal *idsB2* was left intact and we generated two complementation strains (ΔMEV +pMEV_+*idsB2*_ and ΔMEV +pMEV_-*idsB2*_). We found that both ΔMEV and ΔMEV +pMEV_+*idsB2*_ displayed WT growth kinetics with no significant difference in doubling time (Figure 3C,D). However, ΔMEV +pMEV_-*idsB2*_ had a longer doubling time and elongated log phase compared to all the other strains (Figure 3C,D). Further, ΔMEV +pMEV_-*idsB2*_ also formed smaller, more compact colonies that required 24-48 hr of extra incubation to clearly visualize (Supplementary Figure 1B).

### Both pathways are functional and interact at the metabolic level

The ability to delete MEV genes with no obvious defect prompted us to further interrogate the functionality of the MEV pathway. To directly measure intermediates and assess the contribution of the MEV pathway to prenyl phosphate metabolism, we performed metabolic profiling across growth phases (Figure 4A). We observed that the ΔMEV deletion strain had significantly higher levels of acetyl-CoA compared to WT and both complementation strains, all of which encode the MEV pathway (Figure 4B). Together, these data strongly support that the MEV pathway is functional and drawing on the cellular acetyl-CoA pool.

**Figure 4:**
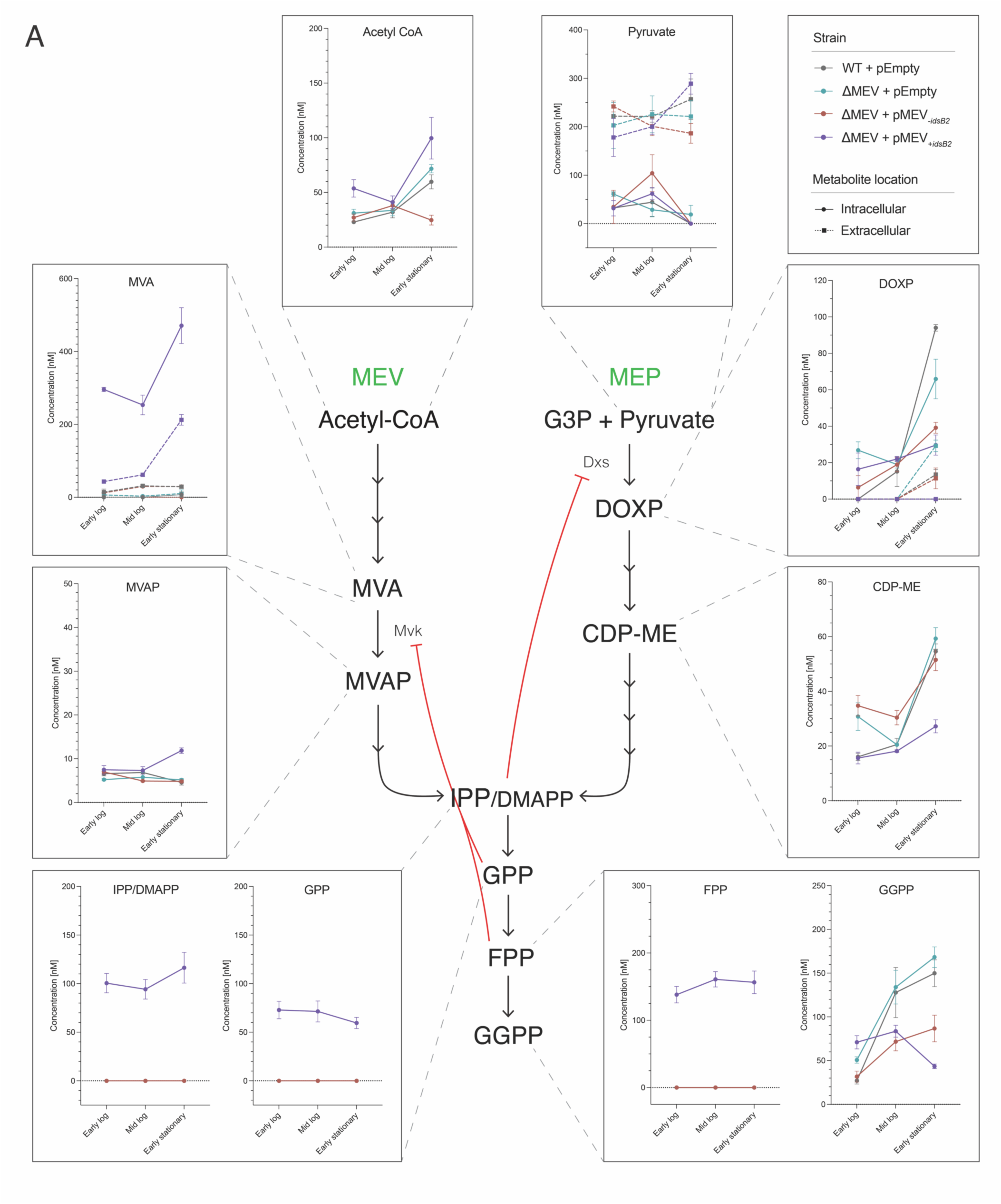
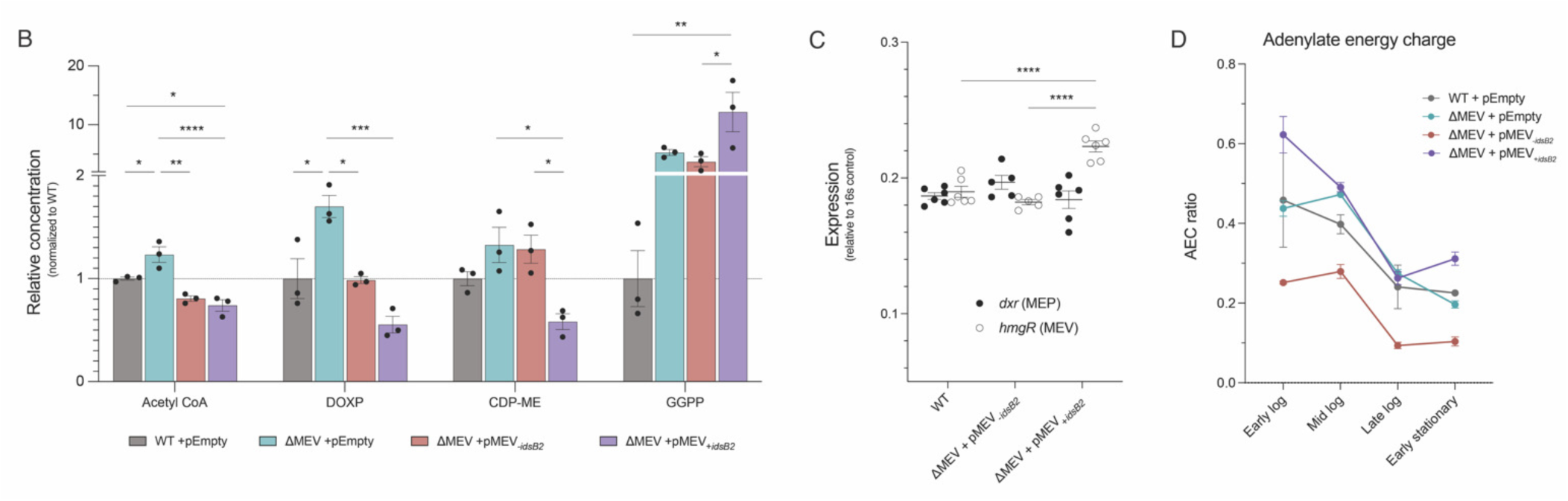
Modulating either isoprenoid biosynthesis pathway impacts metabolism in the other. **A. Metabolic flux of both isoprenoid biosynthesis pathways.** Shown are starting reagents and intermediates of the MEV pathway, the MEP pathway, and polyprenyl metabolism downstream of both pathways at early logarithmic, mid logarithmic, or early stationary phase (OD_600_ 0.2, 0.5, and 2, respectively). Solid lines indicate intracellular metabolites; dotted lines indicate extracellular metabolites. N = 3 biological replicates per strain. **B. MEV expression impacts relative quantities of key metabolites.** ΔMEV has significantly higher levels of acetyl-CoA, DOXP, CDP-ME compared to WT, while ΔMEV +pMEV_+*idsB2*_ has significantly lower levels of these metabolites. Dotted line indicates WT levels. Strains were grown to late logarithmic phase (OD_600_ 1) prior to analysis. Data were analyzed via one-way ANOVA followed by multiple comparisons. *p≤0.05; **p≤0.01; ***p≤0.001; ****p≤0.0001. N = 3 biological replicates per strain. **C. Relative expression of *dxr* and *hmgR* in complemented strains.** ΔMEV +pMEV_+*idsB2*_ had significantly elevated expression of *hmgR*, the rate-limiting step of the MEV pathway (open circles; p<0.0001). Both of the complemented strains had WT expression of *dxr*, representing the MEP pathway (closed circles; p=n.s.). Data shown are relative to 16s mRNA. N ≥ 5 per strain. Data were analyzed by using a two-way ANOVA followed by Dunnett’s multiple comparisons test. **D. Adenylate energy charge (AEC) ratio to evaluate cellular energetics.** AEC ratio is significantly lower in ΔMEV +pMEV_-*idsB2*_ compared to all other strains, reflecting a low energy state in this strain across growth phases. N = 3 biological replicates per strain.

We detected significant accumulation of mevalonate (MVA) and mevalonate-5-phosphate (MVAP) in ΔMEV +pMEV_+*idsB2*_ (Figure 4A), suggesting that this strain has dysregulated flux through the MEV pathway. We did not detect MEV metabolites in the other strains, likely because intermediates of isoprenoid precursor biosynthesis and downstream prenyl pyrophosphate metabolism are normally maintained at very low levels in the cell due to efficient turnover. This dysregulation is also reflected in the expression of *hmgR*, which is significantly elevated in ΔMEV +pMEV_+*idsB2*_ compared to either ΔMEV +pMEV_-*idsB2*_ or WT (Figure 4C). This strain also had significantly lower levels of acetyl-CoA compared to WT (Figure 4B), demonstrating that MEV overexpression depletes starting substrates of the pathway.

Intracellular levels of isoprenoid intermediates are tightly regulated in both pathways (16, 52–54); thus our manipulation of the MEV pathway may impact isoprenoid flux via feedback regulation. We measured polyprenyl intermediates downstream of both the MEV and MEP pathways and found that only ΔMEV +pMEV_+*idsB2*_ had detectable levels of IPP/DMAPP, geranyl pyrophosphate (GPP), and farnesyl pyrophosphate (FPP) (Figure 4A), further demonstrating dysregulated isoprenoid flux in this strain. Accumulation of FPP and GPP is known to feedback inhibit the enzyme mevalonate kinase Mvk (55–57), which we indeed observed in this strain as evidenced by significantly lower levels of MVAP compared to MVA in the immediately preceding step (Figure 4A). While geranylgeranyl diphosphate (GGPP) was detected in all strains, it was highest in ΔMEV +pMEV_+*idsB2*_ (Figure 4B), supporting a role for *idsB2* in catalyzing the formation of GGPP.

Our metabolic profiling also demonstrated interplay between the MEV and MEP pathways. The ΔMEV mutant had elevated levels of MEP intermediates, particularly of 1-deoxy-D-xylulose-5-phosphate (DOXP; Figure 4B), while ΔMEV +pMEV_+*idsB2*_ had lower levels of both DOXP and 4-(cytidine 5′-diphospho)-2-C-methyl-erythritol (CDP-ME; Figure 4B). This suggests a “seesaw” effect of modulating either pathway; a deletion in the MEV pathway causes a compensatory increase in MEP flux, and elevated MEV flux suppresses the MEP pathway.

To assess cellular energetics in our strains, we determined the adenylate energy charge (AEC) ratio, which measures cellular concentrations of ATP, ADP, and AMP to assess the energy stored in the cellular adenine nucleotide pool (41). In mycobacteria, reported AEC ratios range from 0.4-0.8 depending on the species and growth condition (58–60). We found that ΔMEV had a similar AEC ratio to WT and ΔMEV +pMEV_+*idsB2*_ (Figure 4D), indicating that deleting the MEV pathway does not significantly impact the energetic state of the cell. Interestingly, we found that the AEC of ΔMEV +pMEV*_-idsB2_* was an average of two-fold lower than WT across growth phases (SD±0.489; Figure 4D), suggesting that the expression of the MEV pathway without *idsB2* induces a low energy state in Mm. Cellular concentrations of dinucleotide cofactors NADH and NADPH were comparable among the strains, indicating similar redox states (Supplementary Figure 1C,D). The NADH:NAD+ ratio was inversely correlated with AEC ratio in all strains (Figure 4D, Supplementary Figure 1C), likely due to NADH oxidation leading to the synthesis of ATP via the electron transport chain.

Cumulatively, these metabolic findings demonstrate that the MEV pathway is functional and draws from cellular acetyl-CoA pools, and that modulating flux of either pathway has metabolic consequences on the other pathway. Further, the differences observed between the complemented strains support a model in which *idsB2* plays a key role in the fate of prenyl phosphate metabolism.

### ΔMEV has a competitive defect, and complementation with MEV_-*idsB2*_ provides a competitive advantage

Given that the MEV pathway is nonessential yet functional, we sought to test conditions in which its deletion may be consequential. Some dual pathway-encoding bacteria employ each pathway differentially in response to stresses which may not be represented in axenic culture, such as oxidative stress and community competition (8, 18). To more sensitively probe the role of the MEV pathway, we directly competed WT and the ΔMEV mutant, which can reveal subtle growth phenotypes that may not be detected in axenic culture. We found that ΔMEV was outcompeted by WT, and that complementation with pMEV_+*idsB2*_ rescues this defect (Figure 5A,B). Interestingly, complementation with pMEV_-*idsB2*_ reverses the competitive defect of ΔMEV, instead resulting in a strong outcompetition of WT by ΔMEV +pMEV*_-idsB2_* (Figure 5C). This competitive advantage is intriguing because ΔMEV +pMEV_-*idsB2*_ has a longer doubling time and lower AEC ratio. These competitive phenotypes remained consistent under hypoxic growth (Supplementary Figure 2A-C), suggesting that the presence of *idsB2* in the plasmid, rather than the environmental condition, is responsible for the observed competitive phenotypes. To address whether competition was due to a secreted factor, we assessed growth of WT Mm in spent media from either WT, ΔMEV +pMEV_-*idsB2*_, or a co-culture of the two strains. We found no significant differences in WT growth among the spent media (Figure 5D), suggesting that ΔMEV +pMEV_-*idsB2*_ is not secreting a stable factor that restricts the growth of competing bacteria. These data highlight a potential role for *idsB2* in steering the direction of downstream isoprenoid biosynthesis, the end products of which may influence success in a competitive environment.

**Figure 5:**
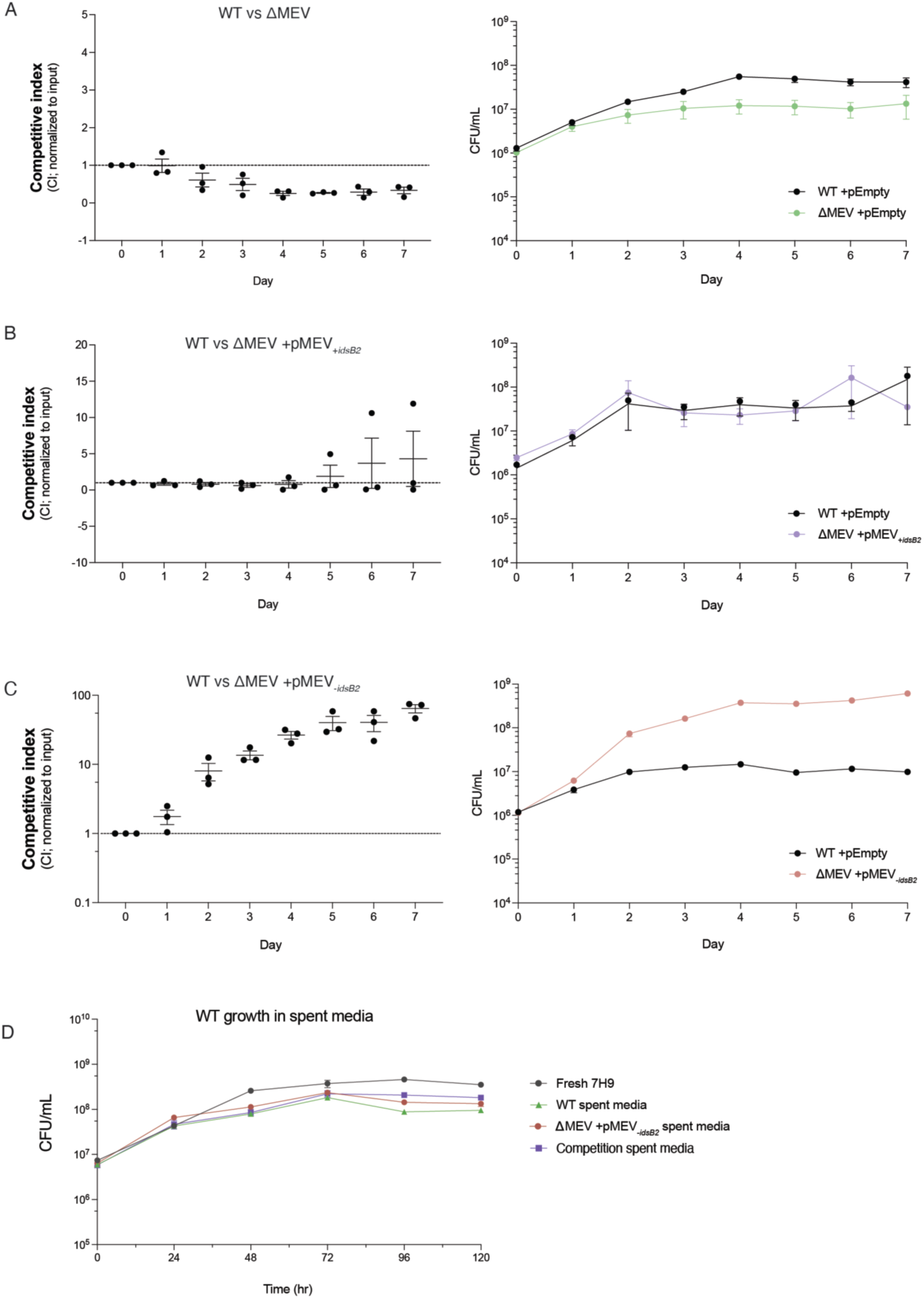
ΔMEV has a competitive defect compared to WT, but ΔMEV +pMEV_-*idsB2*_ outcompetes WT. **A-C. Competitive phenotypes of all strains compared to WT.** WT, ΔMEV, and complemented strains were directly competed, revealing a competitive defect in ΔMEV (**A**) which can be rescued by complementation with pMEV_+*idsB2*_ (**B**). Complementation with pMEV_-*idsB2*_ resulted in strains that strongly outcompeted WT (**C**). Competitive index (CI) of 1 indicates strains performed comparably; CI>1 indicates mutant performed better than WT; CI<1 indicates WT performed better. pEmpty indicates empty vector control. **D. Outcompetition of ΔMEV +pMEV_-*idsB2*_ over WT is not due to secreted factors.** There are no notable differences in WT growth between spent media from axenic or competition cultures. ‘Competition spent media’ indicates spent media isolated from WT vs ΔMEV +pMEV_-*idsB2*_ competition cultures.

### The MEV pathway is dispensable during infection

To test whether the MEV pathway plays a role in survival of host immunity, we tested the ΔMEV strain using two widely-used models of infection: *ex vivo* macrophage and *in vivo* zebrafish infections (61). We found that survival of Mm in murine bone marrow-derived macrophages (BMMs) is not significantly different among the strains, as the ΔMEV mutant grew with WT kinetics in both resting and interferon gamma (IFNγ)-activated macrophages (Figure 6A,B). Further, there were no significant differences in macrophage cell death, underscoring the similarity of the strains during infection (Supplementary Figure 3A,B). We then turned to the well-established larval zebrafish infection model to assess any strain differences and observed similar *in vivo* replication between WT and ΔMEV (Figure 6C). Additionally, we found that both strains were able to disseminate to the brain in roughly half of the infections, indicating no overall difference in ability to colonize the host (Figure 6D). Together, these infection data support that the MEV pathway does not play a significant role in growth and survival *in vivo*.

**Figure 6:**
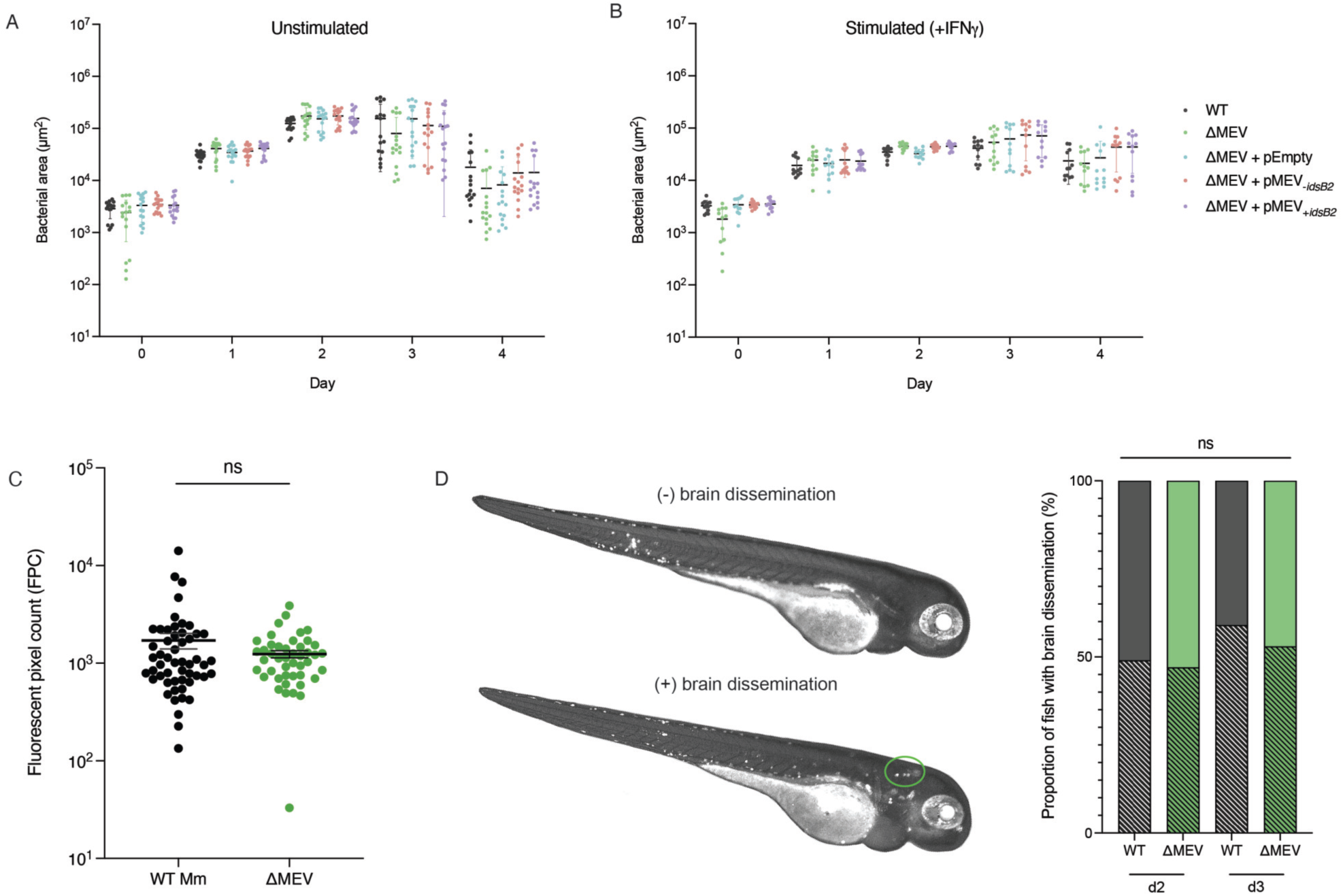
The MEV pathway does not contribute to survival during infection. **A,B. Growth and survival in macrophages is not significantly different between the strains.** Wild-type BMMs were plated and grown with (**A**) or without (**B**) the activating cytokine IFNγ and subsequently infected at an MOI of 1 with various strains of Mm. For four days following infections, cells were fixed and imaged and the bacterial fluorescence and nuclei count were measured. Data plotted are the mean area of bacterial fluorescence ± SEM. N = 3 biological replicates. **C,D. ΔMEV exhibits WT growth and dissemination to the brain in zebrafish.** There is no significant difference in *in vivo* growth (**C**) or dissemination to the brain (**D**) between WT and ΔMEV strains. Larval zebrafish were infected with various strains of Mm via caudal vein injection, and fluorescent pixel count (FPC) was measured on day 2 post infection. Error bars represent mean FPC ± SEM (**C**; p=0.1803; N = 3 infections) or percent of disseminated bacteria (**D**; p=0.3471; N = 3 infections). FPC comparisons were tested with an unpaired t-test and dissemination comparisons were tested with a Fisher’s exact test.

### MEV expression has functional consequences under environmental stresses

Evidence from *Listeria* suggests that the MEP and MEV pathways may be used differently in dual pathway-encoding bacteria based on oxygen availability (18). To address whether this is true in Mm, we used the Wayne model of nonreplicating persistence (43) to induce hypoxia in our strains. We observed no difference in OD_600_ or CFU among the strains throughout hypoxia (Supplementary Figure 4A), although we did observe a qualitative difference in oxygen utilization among the strains using the oxygen indicator methylene blue (Supplementary Figure 4B), prompting us to more closely assess strain differences under hypoxia. We measured pathway expression during hypoxic adaptation and found significant overexpression of *hmgR*, the rate-limiting step of the MEV pathway, early in hypoxia compared to *dxr*, particularly on day 4 (Figure 7A). Further, published RNAseq data have identified that several MEV genes are upregulated in Mm during hypoxia and shortly after re-aeration, including *pksG, hmgR*, *idsB1*, *mpd*, and *erg12* (62, 63). Together, these data suggest that utilization of the MEV pathway may be linked to oxygen availability.

**Figure 7:**
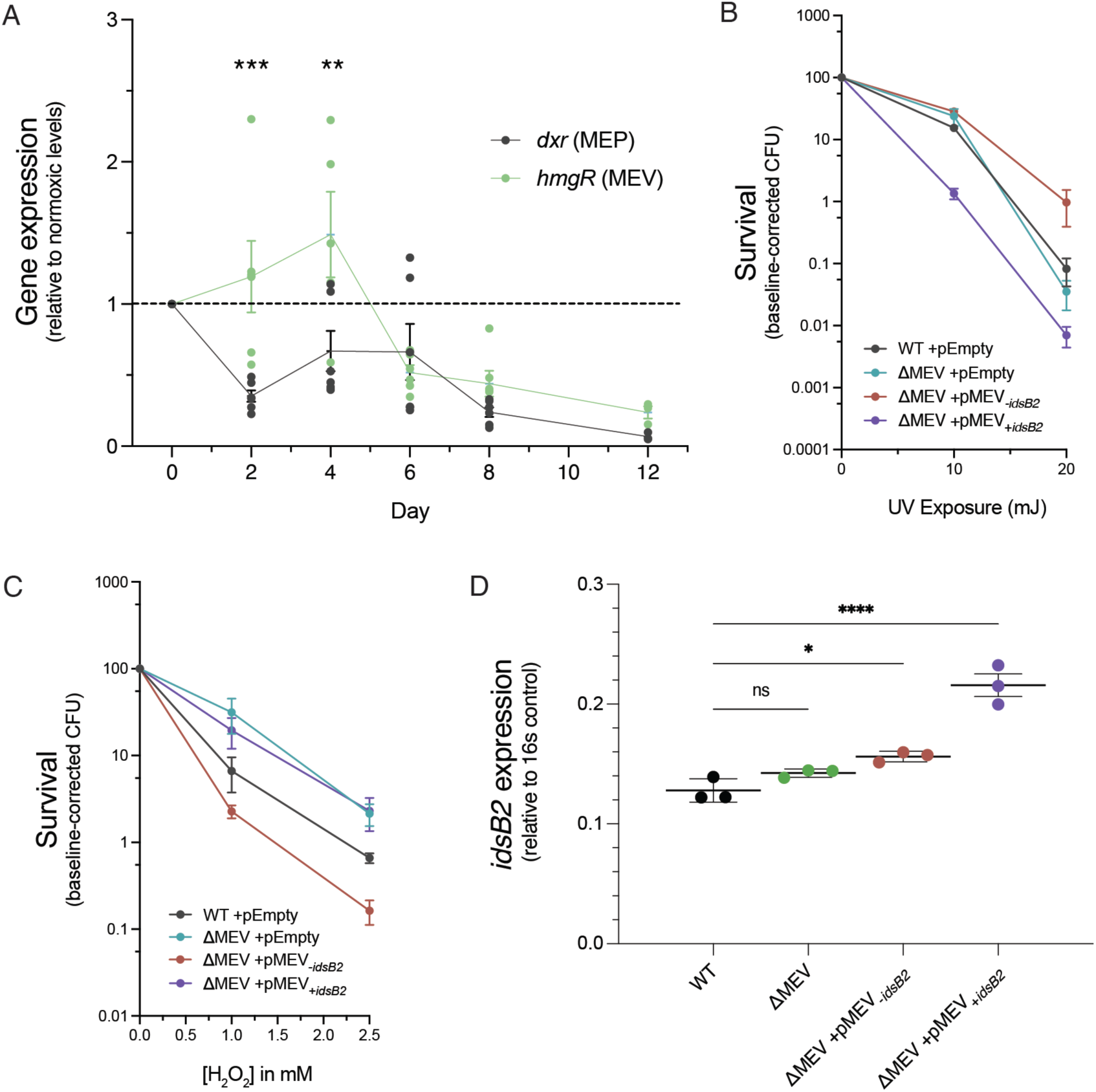
MEV pathway expression has functional consequences under environmental stresses. **A. The MEV pathway is significantly upregulated early in the shift down to hypoxia.** There is a significant difference in overall pathway expression between timepoints (two-way ANOVA; p=0.0006) and on days 2 and 4 of hypoxia, *hmgR* was significantly overexpressed compared to *dxr* (Sidak’s multiple comparisons; p=0.0004 and 0.0010 respectively). N=6 biological replicates. **B,C. Functional consequences of modulating MEV.** ΔMEV has WT resistance to UV damage (**B**) but is more resistant to H_2_O_2_ (**C**). Complemented strains are differentially sensitive to UV (**B**) and H_2_O_2_ stress (**C**). Strains were grown to mid-log, OD-matched, diluted, and exposed to UV light doses ranging from 0 to 20 mJ (**B**) or to H_2_O_2_ concentrations ranging from 0 to 2.5 mM (**C**). pEmpty indicates empty vector control. **D. Relative expression of *idsB2* in all strains.** The knockout ΔMEV had WT levels of *idsB2* expression (p=n.s.) while ΔMEV +pMEV_-*idsB2*_ had slightly elevated expression (p=0.0202), potentially due to the reintroduction of regulatory elements on the plasmid. ΔMEV +pMEV_+*idsB2*_ had significantly more *idsB2* expression compared to WT (p<0.0001). Data shown are relative to 16s mRNA. N = 3 per strain. Data were analyzed using one-way ANOVA followed by Tukey’s multiple comparisons.

Mm is thought to protect itself from lethal photooxidation by producing β-carotene, a photoinducible isoprenoid that gives Mm its orange pigment (64–66). We observed WT levels of β-carotene in ΔMEV (Supplementary Figure 4C), suggesting that the MEV pathway does not play a critical role in its synthesis. However, we found that the complementation strains produced less β-carotene compared to WT and ΔMEV (Supplementary Figure 4C), suggesting that multi-copy expression of the MEV pathway has a functional consequence on isoprenoid end products. To determine how differing levels of β-carotene may impact resistance to photooxidation from UV exposure, we exposed the strains to varying levels of UV and assayed for survival. We found no differences between WT and ΔMEV, in agreement with our observations of the strains producing equivalent levels of β-carotene; however, the two complemented strains were differentially sensitive to UV damage (Figure 7B). These data suggest a role for *idsB2* copy number in determining the fate of isoprenoids, particularly of the β-carotene pigment.

To test whether the MEV pathway plays a role in survival of oxidative challenge, we exposed our strains to hydrogen peroxide (H_2_O_2_) and assessed survival after six hours. We found that both ΔMEV and ΔMEV +pMEV_+*idsB2*_ were more resistant to peroxide stress, while ΔMEV +pMEV_-*idsB2*_ was more sensitive compared to WT (Figure 7C). This might be explained by an increase in the production of the oxidative stress signaling molecule MEcPP in ΔMEV, which has elevated flux through the MEP pathway (Figure 4). As we observed with UV exposure, the complementation strains were differentially susceptible to H_2_O_2_, suggesting *idsB2* directs isoprenoid synthesis toward certain end products that may be critical to surviving disparate environmental stresses. We observed that while ΔMEV had WT expression of *idsB2*, ΔMEV +pMEV_+*idsB2*_ had significantly higher expression compared to WT, demonstrating that *idsB2* on the multicopy plasmid indeed results in more mRNA (Figure 7D). Taken together, these findings suggest that regulation of the MEV pathway supports flexibility of Mm under variable environmental challenges, including hypoxic, oxidative, and UV stress. Further, it appears that copy number of *idsB2* is important to direct downstream prenyl phosphate metabolism, and that the observed phenotypes among the strains may be driven by the identity of isoprenoid end products.

## Discussion

In this study, we investigated the MEP and the MEV pathways of isoprenoid biosynthesis to probe the evolutionary pressures that led to Mm encoding both. Using a reverse genetics approach, we manipulated either pathway to assess the contribution of each to growth and physiology, presenting the first study to investigate the functional role of both isoprenoid biosynthesis pathways in Mm. Prior to this study, it was unknown whether these pathways were redundant in Mm, which is one of the few mycobacterial species that encodes both the MEV and MEP pathways. We found that the MEP pathway is essential in Mm, providing critical functional validation of prior high-throughput essentiality experiments which failed to identify several MEP genes as essential (23, 24). Further, we showed that the MEV pathway is nonessential and used targeted metabolomics to demonstrate that the MEV pathway is functional. Importantly, we found metabolic interplay between the two pathways in Mm.

Not only is MEV pathway usage consequential in certain conditions, but our data further suggest that *idsB2* plays an important role in MEV-derived isoprenoid biosynthesis. We found that dosage of the *idsB2* gene had functional consequences on our strains with variable MEV flux, which were differentially susceptible to UV- and H_2_O_2_-mediated killing. Further, ΔMEV +pMEV_-*idsB2*_ has a small colony phenotype and growth defect, consistent with its lower energetic state. IdsB2 is a polyprenyl synthetase whose exact function has not been experimentally demonstrated in Mm, although it is likely involved in the synthesis of a MEV-dependent isoprenoid due to its inclusion in the MEV operon. IdsB2 shares high amino acid (57.06% identity) and structural (0.822 Å root mean squared) homology with IdsB in *M. tuberculosis* (Supplementary Figure 4D, Supplementary Table 5), which has been shown to catalyze the condensation of IPP with E,E-FPP to form GGPP (67). Mm encodes four IdsB homologs, likely due to functional diversification; thus, the specific role of IdsB2 in downstream isoprenoid metabolism remains to be experimentally determined. However, because of high sequence and structural homology, we suspect an analogous function for IdsB2 in Mm. One possible explanation for why *idsB2* is encoded in the MEV operon may be to promote GGPP synthesis from FPP under conditions in which the MEV pathway is upregulated, thus relieving feedback inhibition of Mvk by FPP. Further, perhaps the MEV operon contains *idsB2* to accumulate GGPP and feedback inhibit the MEP pathway to more fully switch to MEV synthesis under certain conditions, although no feedback inhibition of the MEP pathway by GGPP has been reported to date.

Further, stress conditions may impact substrate availability, which may underlie differential pathway usage in this system. Each pathway uses different starting substrates, and certain stresses may perturb homeostatic metabolism such that using one pathway over the other is favoured. A putative sigma factor F (SigF) binding motif can be found in the promoter region of the MEV operon, suggesting that the MEV pathway might indeed play a role in the stress response of Mm by expanding its metabolic repertoire. In support of such a model, SigF has been shown to regulate both carotenoid production and resistance to H_2_O_2_ in *M. smegmatis* (69). While the SigF regulons have been defined in *M. tuberculosis* and *M. smegmatis* (70, 71), these organisms do not encode the MEV pathway so no functional information is available for the potential SigF regulation of the MEV operon at this time. Regulation of the MEV operon may not only provide a different means to generate IPP/DMAPP under stress but also influences the identity of mature isoprenoids by influencing downstream enzymatic steps. Taken together, these data support a model that the MEV pathway provides metabolic flexibility in Mm.

Evidence from previous bioengineering work further supports a model of the MEV pathway affording Mm metabolic flexibility. Engineering efforts in *E. coli* have demonstrated synergy between the two pathways (72), suggesting that perhaps dual encoding bacteria like Mm harness this synergy to generate a higher yield of critical isoprenoids than could be achieved by a single pathway. Alternatively, dual encoding bacteria may use the MEV pathway as a failsafe to ensure isoprenoid flux continues in the case of MEP blockage or prenyl phosphate intermediate accumulation. In addition to the presence of MEV as a metabolic failsafe, our data suggest that *idsB2* may serve as yet another layer of control over end product fate. Given the divergent metabolism and phenotypes of the two complementation strains, which differ only in the copy number of *idsB2*, it appears that the level of *idsB2* expression may serve as a determinant of the biosynthetic fate of both MEV- and MEP-derived metabolites. These findings highlight a contrast between the role of the MEV pathway in Mm and other bacteria.

Recently the MEP pathway has emerged as an oxidative stress signaling pathway (16), with striking evidence for this role in *Listeria* species that encode both pathways of isoprenoid biosynthesis. However, these *Listeria* primarily utilize the MEV pathway for IPP synthesis and encode incomplete MEP pathways. In contrast, Mm uses the MEP pathway for essential isoprenoid metabolism and encodes both pathways fully intact. Thus, it appears that these pathways play different roles in *Listeria* and Mm: oxidative stress sensing in *Listeria* and metabolic flexibility in Mm.

Our work provides novel and valuable insights about the interactions between both pathways at the functional and metabolic levels. Our data support that the essential MEP pathway facilitates central metabolism, while the dispensable MEV pathway may represent an inducible accessory system that supports metabolic flexibility in dynamic environmental niches. Further, we demonstrate for the first time that modulation of flux in either pathway causes a compensatory shift in the other, suggesting that the pathways are functional and interacting at the metabolic level. Such insights are critical to understand how Mm leverages the MEV pathway and further broadens our ability to engineer its close mycobacterial relatives, like the important human pathogens *M. tuberculosis* and *M. avium*.

## Acknowledgements

We thank Zoila Álvarez-Aponte, Hans Carlson, Kathleen Ryan, Sarah Stanley, Rebecca Procknow, Aziz Qabar, Astrid, and all members of the Cox lab for their helpful discussions. This work was supported by the National Institutes of Health grants T32GM132022 (C.M.Q.), U19AI162583 (J.S.C.), U19AI135990 (J.S.C.), 1P01 AI063302 (D.A.P.), 1R01 AI027655 (D.A.P.), 5R01 CA283604 (D.A.P.), 1DP2NS127277 (C.A.M. and T.Q.), and the Henry Wheeler Center for Emerging and Neglected Diseases (C.M.Q.). This material was based upon work supported by the Joint BioEnergy Institute, U.S. Department of Energy, Office of Science, Biological and Environmental Research Program under Award Number DE-AC02-05CH11231 with Lawrence Berkeley National Laboratory.

## Supplementary figure legends

**Figure S1:**
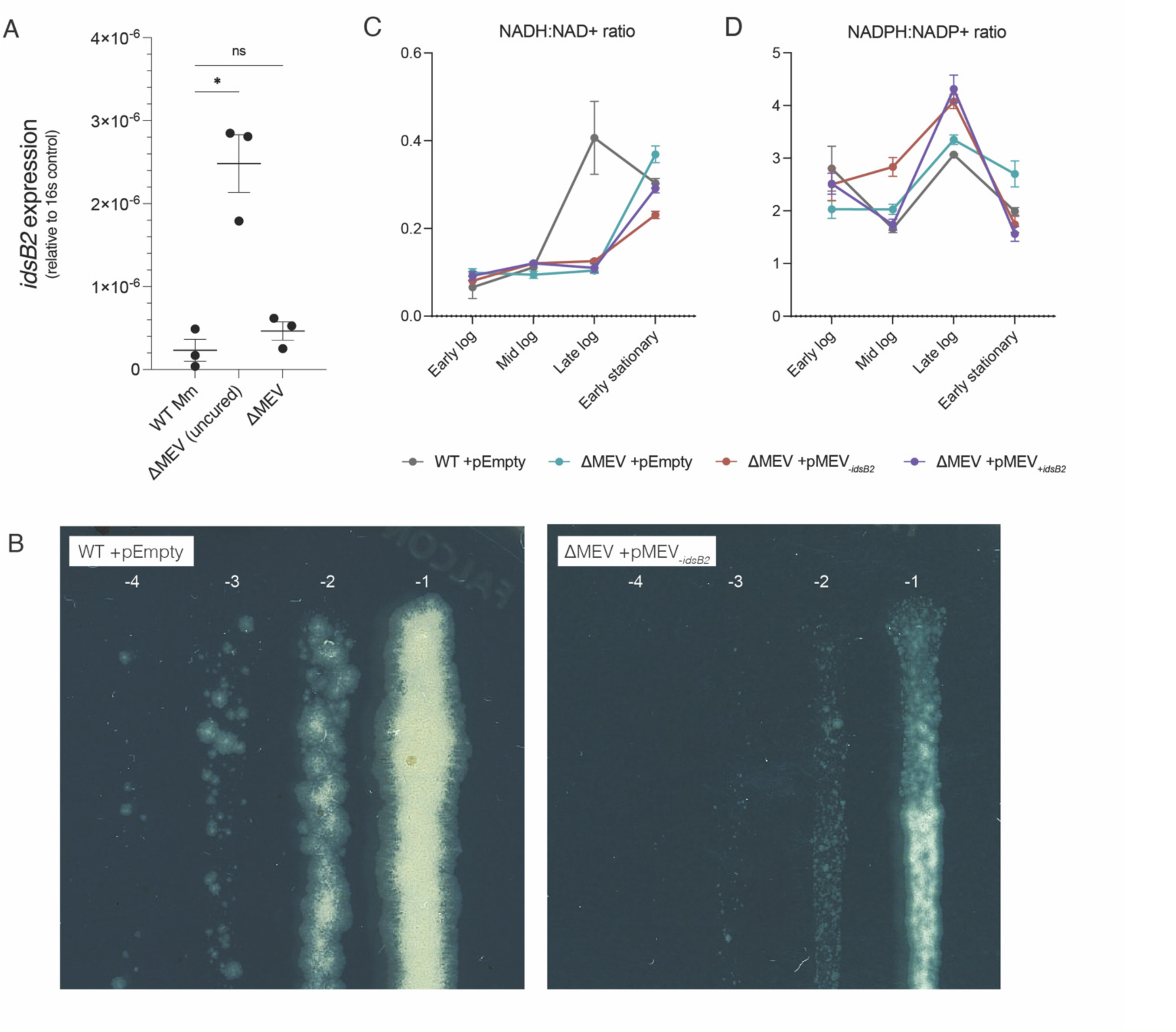
Initial strain characterization. **A. Relative expression of *idsB2* in WT, ORBIT-integrated ΔMEV, and cured ΔMEV.** Integration of the ORBIT plasmid to generate the ΔMEV knockout strain caused strong polar effects on the expression of the downstream gene *idsB2* (p=0.0208). Upon excising the integrated plasmid from this strain, *idsB2* levels returned to WT (p=n.s.). Data shown are relative expression of *idsB2* compared to 16s mRNA. N = 3 per strain. Data were analyzed by fitting a mixed-effects model followed by Dunnett’s multiple comparisons test. **B. ΔMEV +pMEV_-*idsB2*_ forms small colonies.** Mid-log cultures of WT and ΔMEV +pMEV_-*idsB2*_ were serially diluted and plated on 7H10 agar. Plates were incubated at 30°C for five days prior to imaging. **D,E. Dinucleotide cofactors to assess cellular redox states.** NADH:NAD+ **(D)** and NADPH:NADP+ **(E)** ratios across growth phases. NADH:NAD+ ratio is inversely correlated with AEC ratio. N = 3 biological replicates per strain.

**Figure S2:**
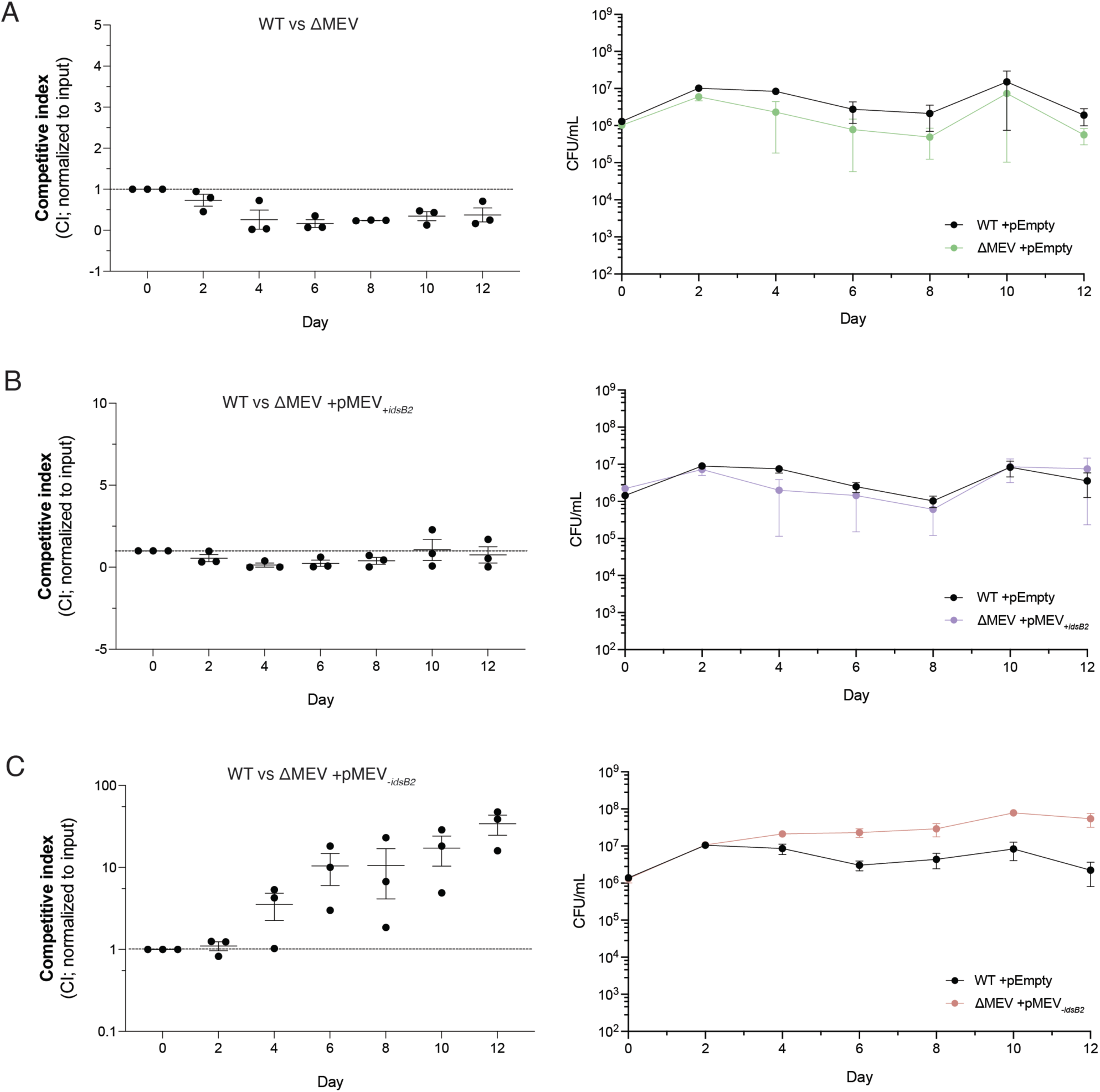
Similarly to normoxic competition, ΔMEV has a competitive defect while ΔMEV +pMEV_-*idsB2*_ outcompetes WT. The strains were directly competed as in figure 5 (normoxic competition) but in sealed tubes per the Wayne model. As observed in normoxia, ΔMEV had a competitive defect (**A**) which can be rescued by complementation with MEV_+*idsB2*_ (**B**), but complementation with MEV_-*idsB2*_ strongly outcompeted WT (**C**). Cultures were plated every other day for twelve days on selective media. CFU were counted to calculate competitive index (CI) normalized to input.

**Figure S3:**
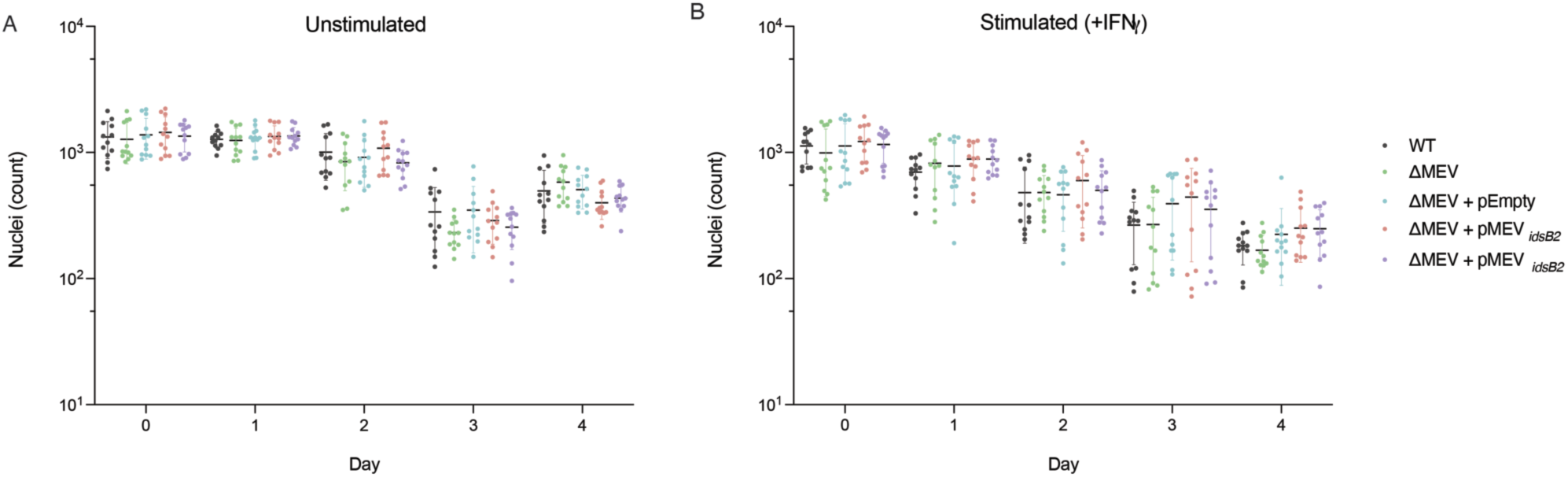
Macrophage cell death. **Macrophage cell death is not significantly different between the strains.** Wild-type bone marrow-derived macrophages (BMMs) were infected at an MOI of 1 as previously described, either without (**A**) or with (**B**) IFNγ. Plates were imaged on a PerkinElmer OperaPhenix confocal microscope and number of nuclei were quantified via the Harmony software. Data plotted are mean nuclei count ± SEM. N=3 biological replicates; 4 technical replicates per biological replicate.

**Figure S4:**
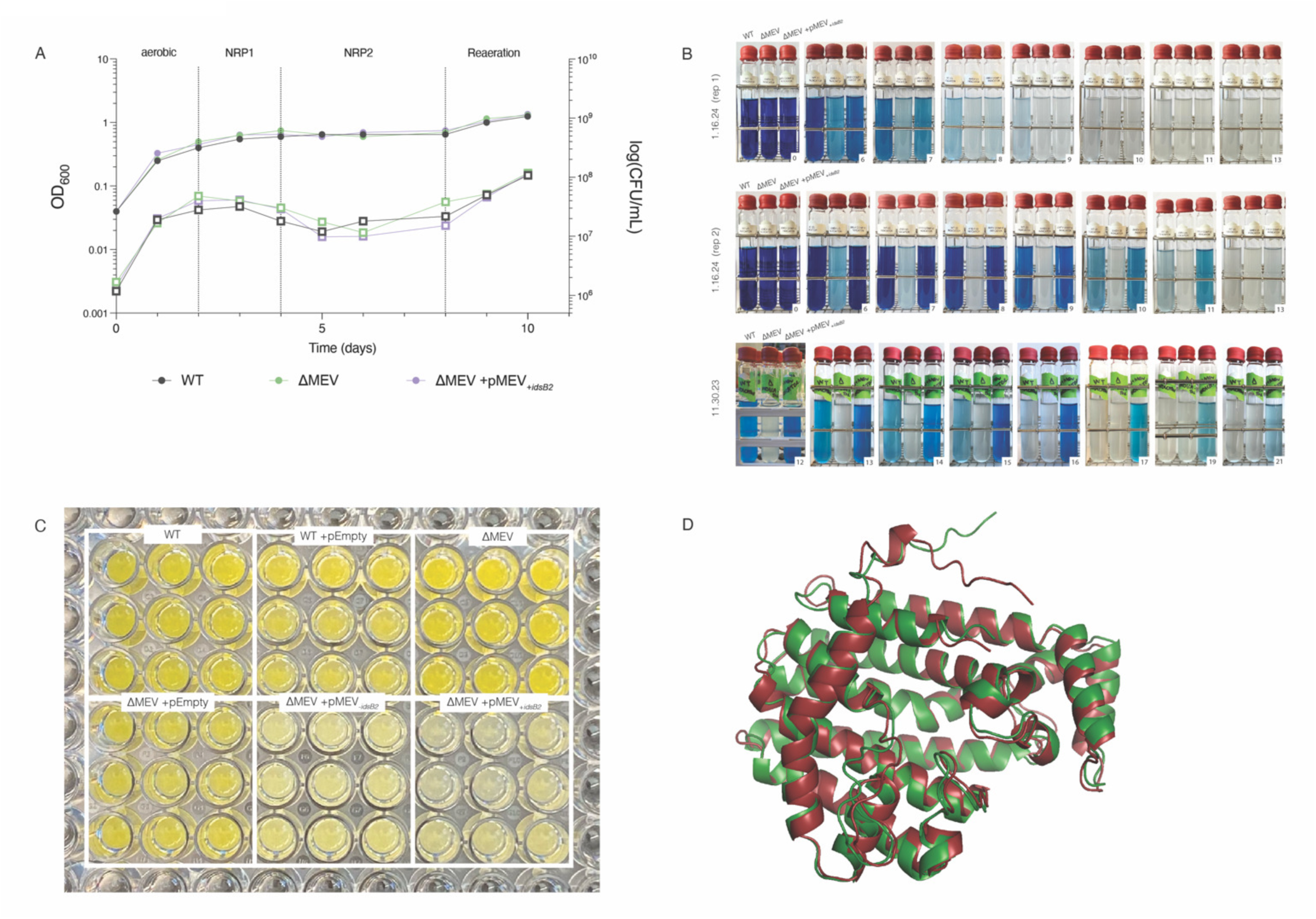
Environmental stress and response. **A. There is no significant difference in OD or CFU throughout hypoxia among the strains.** Strains were slowly induced for hypoxia using the Wayne model. Cultures were started at OD_600_ 0.05 in glass tubes with 8mm stir bars to a headspace ratio of 0.5. Tubes were sealed with a rubber stopper and incubated at 30°C with stirring. OD_600_ and CFU samples were taken daily for a minimum of 10 days. To avoid reaeration during the time course, one tube was designated per timepoint. **B. MmΔMEV decolorizes quickly in hypoxia, while the complemented strain decolourizes more slowly compared to WT.** WT Mm (left), ΔMEV (middle), and ΔMEV +pMEV_+*idsB2*_ (right) were slowly induced for hypoxia as described above. Aeration of the cultures was visualized with the addition of methylene blue (25 µg/mL). Three biological replicates are shown by row. Days after induction of hypoxia are labeled in the bottom right corner. **C**. **Complementation strains produce less β-carotene over eight days of growth.** Strains were grown for eight days and OD_600_ was measured to generate the growth curve in Figure 3C. Following this period, the growth plate was left at room temperature in light for 2 days to develop prior to imaging. **D**. **IdsB2 has high predicted structural homology to IdsB in *M. tuberculosis*.** Shown are the aligned predicted structures of Mm IdsB2 (green) and *M. tuberculosis* IdsB (red). Root mean squared (RMSD) is 0.822 Å.

## Supporting information

### Supplementary Tables

Table S1: 16s rRNA sequences used to generate phylogenetic tree.

Table S2: Primers and oligos used in this study.

Table S3: Plasmids used in this study.

Table S4: Protein sequences used to generate hmgR alignment.

Table S5: Distance matrix of IdsB homologs. Shown is percent identity of IdsB homologs in Mm and *M. tuberculosis* (Mtb).

### Supplementary Files

File S1: MUSCLE alignment of 84 mycobacterial 16s rRNA sequences.

File S2: MUSCLE alignment of amino acid sequences of class I (*Mycobacterium marinum, Actinoplanes sp.* A40644, *Vibrio cholerae, Streptomyces rutgerensis, Nocardia brasiliensis, Corynebacterium striatum*) and class II (*Listeria monocytogenes, Streptococcus pneumoniae, Staphylococcus aureus, Rhodococcus erythropolis,* and *Pseudomonas mevalonii*) HmgR enzymes.

File S3: Raw metabolomics data.

## References

1. Faylo JL, Ronnebaum TA, Christianson DW. 2021. Assembly-Line Catalysis in Bifunctional Terpene Synthases. Acc Chem Res 54:3780–3791.

2. Sacchettini JC, Poulter CD. 1997. Creating Isoprenoid Diversity. Science 277:1788–1789.

3. Avalos M, Garbeva P, Vader L, Wezel GP van, Dickschat JS, Ulanova D. 2021. Biosynthesis, evolution and ecology of microbial terpenoids. Nat Prod Rep 39:249–272.

4. Eberl M, Hintz M, Reichenberg A, Kollas A-K, Wiesner J, Jomaa H. 2003. Microbial isoprenoid biosynthesis and human γδ T cell activation. Febs Lett 544:4–10.

5. Heuston S, Begley M, Gahan CGM, Hill C. 2012. Isoprenoid biosynthesis in bacterial pathogens. Microbiology+ 158:1389–1401.

6. Ahyong V, Berdan CA, Burke TP, Nomura DK, Welch MD. 2019. A Metabolic Dependency for Host Isoprenoids in the Obligate *In*tracellular Pathogen Rickettsia parkeri Underlies a Sensitivity to the Statin Class of Host-Targeted Therapeutics. mSphere 4:e00536–19.

7. Boucher Y, Douady CJ, Papke RT, Walsh DA, Boudreau MER, Nesbø CL, Case RJ, Doolittle WF. 2003. Lateral gene transfer and the origins of prokaryotic groups. Annu Rev Genet 37:283–328.

8. Boucher Y, Doolittle WF. 2000. The role of lateral gene transfer in the evolution of isoprenoid biosynthesis pathways. Mol Microbiol 37:703–716.

9. Lombard J, Moreira D. 2010. Origins and Early Evolution of the Mevalonate Pathway of Isoprenoid Biosynthesis in the Three Domains of Life. Mol Biol Evol 28:87–99.

10. Stinear TP, Seemann T, Harrison PF, Jenkin GA, Davies JK, Johnson PDR, Abdellah Z, Arrowsmith C, Chillingworth T, Churcher C, Clarke K, Cronin A, Davis P, Goodhead I, Holroyd N, Jagels K, Lord A, Moule S, Mungall K, Norbertczak H, Quail MA, Rabbinowitsch E, Walker D, White B, Whitehead S, Small PLC, Brosch R, Ramakrishnan L, Fischbach MA, Parkhill J, Cole ST. 2008. Insights from the complete genome sequence of Mycobacterium marinum on the evolution of Mycobacterium tuberculosis. Genome Res 18:729–741.

11. Altincicek B, Moll J, Campos N, Foerster G, Beck E, Hoeffler J-F, Grosdemange-Billiard C, Rodríguez-Concepción M, Rohmer M, Boronat A, Eberl M, Jomaa H. 2001. Cutting Edge: Human γδ T Cells Are Activated by Intermediates of the 2-C-methyl-d-erythritol 4-phosphate Pathway of Isoprenoid Biosynthesis. J Immunol 166:3655–3658.

12. Hintz M, Reichenberg A, Altincicek B, Bahr U, Gschwind RM, Kollas A-K, Beck E, Wiesner J, Eberl M, Jomaa H. 2001. Identification of (E)-4-hydroxy-3-methyl-but-2-enyl pyrophosphate as a major activator for human γδ T cells in Escherichia coli. FEBS Lett 509:317–322.

13. Tanaka Y, Morita CT, Tanaka Y, Nieves E, Brenner MB, Bloom BR. 1995. Natural and synthetic non-peptide antigens recognized by human gamma delta T cells. Nature 375.

14. Adam P, Hecht S, Eisenreich W, Kaiser J, Gräwert T, Arigoni D, Bacher A, Rohdich F. 2002. Biosynthesis of terpenes: Studies on 1-hydroxy-2-methyl-2-(E)-butenyl 4-diphosphate reductase. Proc Natl Acad Sci 99:12108–12113.

15. Wolff M, Seemann M, Grosdemange-Billiard C, Tritsch D, Campos N, Rodríguez-Concepción M, Boronat A, Rohmer M. 2002. Isoprenoid biosynthesis via the methylerythritol phosphate pathway. (E)-4-Hydroxy-3-methylbut-2-enyl diphosphate: chemical synthesis and formation from methylerythritol cyclodiphosphate by a cell-free system from Escherichia coli. Tetrahedron Lett 43:2555–2559.

16. Perez-Gil J, Behrendorff J, Douw A, Vickers CE. 2024. The methylerythritol phosphate pathway as an oxidative stress sense and response system. Nat Commun 15:5303.

17. Ostrovsky D, Diomina G, Lysak E, Matveeva E, Ogrel O, Trutko S. 1998. Effect of oxidative stress on the biosynthesis of 2-C-methyl-d-erythritol-2,4-cyclopyrophosphate and isoprenoids by several bacterial strains. Arch Microbiol 171:69–72.

18. Lee ED, Navas KI, Portnoy DA. 2019. The Nonmevalonate Pathway of Isoprenoid Biosynthesis Supports Anaerobic Growth of Listeria monocytogenes. Infect Immun 88.

19. Begley M, Bron PA, Heuston S, Casey PG, Englert N, Wiesner J, Jomaa H, Gahan CGM, Hill C. 2008. Analysis of the Isoprenoid Biosynthesis Pathways in Listeria monocytogenes Reveals a Role for the Alternative 2-C-Methyl-d-Erythritol 4-Phosphate Pathway in Murine Infection▿. Infect Immun 76:5392–5401.

20. Mains DR, Eallonardo SJ, Freitag NE. 2021. Identification of Listeria monocytogenes Genes Contributing to Oxidative Stress Resistance under Conditions Relevant to Host Infection. Infect Immun 89:10.1128/iai.00700-20.

21. Brown AC, Eberl M, Crick DC, Jomaa H, Parish T. 2010. The Nonmevalonate Pathway of Isoprenoid Biosynthesis in Mycobacterium tuberculosis Is Essential and Transcriptionally Regulated by Dxs ▿ †. J Bacteriol 192:2424–2433.

22. Qabar CM, Waldburger L, Keasling JD, Portnoy DA, Cox JS. 2025. Leveraging a synthetic biology approach to enhance BCG-mediated expansion of Vγ9Vδ2 T cells. bioRxiv 2025.05.05.651767.

23. Weerdenburg EM, Abdallah AM, Rangkuti F, Ghany MAE, Otto TD, Adroub SA, Molenaar D, Ummels R, Veen K ter, Stempvoort G van, Sar AM van der, Ali S, Langridge GC, Thomson NR, Pain A, Bitter W. 2015. Genome-Wide Transposon Mutagenesis Indicates that Mycobacterium marinum Customizes Its Virulence Mechanisms for Survival and Replication in Different Hosts. Infect Immun 83:1778–1788.

24. Lefrançois LH, Nitschke J, Wu H, Panis G, Prados J, Butler RE, Mendum TA, Hanna N, Stewart GR, Soldati T. 2024. Temporal genome-wide fitness analysis of Mycobacterium marinum during infection reveals the genetic requirement for virulence and survival in amoebae and microglial cells. mSystems 9:e01326–23.

25. Alvarez-Aponte ZI, Govindaraju AM, Hallberg ZF, Nicolas AM, Green MA, Mok KC, Fonseca-García C, Coleman-Derr D, Brodie EL, Carlson HK, Taga ME. 2024. Phylogenetic distribution and experimental characterization of corrinoid production and dependence in soil bacterial isolates. ISME J 18:wrae068.

26. Karp PD, Billington R, Caspi R, Fulcher CA, Latendresse M, Kothari A, Keseler IM, Krummenacker M, Midford PE, Ong Q, Ong WK, Paley SM, Subhraveti P. 2017. The BioCyc collection of microbial genomes and metabolic pathways. Brief Bioinform 20:1085–1093.

27. Sayers EW, Bolton EE, Brister JR, Canese K, Chan J, Comeau DC, Connor R, Funk K, Kelly C, Kim S, Madej T, Marchler-Bauer A, Lanczycki C, Lathrop S, Lu Z, Thibaud-Nissen F, Murphy T, Phan L, Skripchenko Y, Tse T, Wang J, Williams R, Trawick BW, Pruitt KD, Sherry ST. 2021. Database resources of the national center for biotechnology information. Nucleic Acids Res 50:D20–D26.

28. Edgar RC. 2004. MUSCLE: multiple sequence alignment with high accuracy and high throughput. Nucleic Acids Res 32:1792–1797.

29. Guindon S, Dufayard J-F, Lefort V, Anisimova M, Hordijk W, Gascuel O. 2010. New Algorithms and Methods to Estimate Maximum-Likelihood Phylogenies: Assessing the Performance of PhyML 3.0. Syst Biol 59:307–321.

30. Letunic I, Bork P. 2021. Interactive Tree Of Life (iTOL) v5: an online tool for phylogenetic tree display and annotation. Nucleic Acids Res 49:W293–W296.

31. Price MN, Dehal PS, Arkin AP. 2010. FastTree 2 – Approximately Maximum-Likelihood Trees for Large Alignments. PLoS ONE 5:e9490.

32. Abramson J, Adler J, Dunger J, Evans R, Green T, Pritzel A, Ronneberger O, Willmore L, Ballard AJ, Bambrick J, Bodenstein SW, Evans DA, Hung C-C, O’Neill M, Reiman D, Tunyasuvunakool K, Wu Z, Žemgulytė A, Arvaniti E, Beattie C, Bertolli O, Bridgland A, Cherepanov A, Congreve M, Cowen-Rivers AI, Cowie A, Figurnov M, Fuchs FB, Gladman H, Jain R, Khan YA, Low CMR, Perlin K, Potapenko A, Savy P, Singh S, Stecula A, Thillaisundaram A, Tong C, Yakneen S, Zhong ED, Zielinski M, Žídek A, Bapst V, Kohli P, Jaderberg M, Hassabis D, Jumper JM. 2024. Accurate structure prediction of biomolecular interactions with AlphaFold 3. Nature 630:493–500.

33. Schrödinger L. The PyMOL Molecular Graphics System, Version∼1.8.

34. Murphy KC, Nelson SJ, Nambi S, Papavinasasundaram K, Baer CE, Sassetti CM. 2018. ORBIT: a New Paradigm for Genetic Engineering of Mycobacterial Chromosomes. Mbio 9:e01467–18.

35. Labun K, Montague T, Krause M, Cleuren YT, Tjeldnes H, Valen E. 2019. CHOPCHOP v3: expanding the CRISPR web toolbox beyond genome editing. Nucleic Acids Research W1:W171–174.

36. Rock JM, Hopkins FF, Chavez A, Diallo M, Chase MR, Gerrick ER, Pritchard JR, Church GM, Rubin EJ, Sassetti CM, Schnappinger D, Fortune SM. 2017. Programmable transcriptional repression in mycobacteria using an orthogonal CRISPR interference platform. Nat Microbiol 2:16274.

37. Meijers AS, Troost R, Ummels R, Maaskant J, Speer A, Nejentsev S, Bitter W, Kuijl CP. 2020. Efficient genome editing in pathogenic mycobacteria using Streptococcus thermophilus CRISPR1-Cas9. Tuberculosis 124:101983.

38. Schmittgen TD, Livak KJ. 2008. Analyzing real-time PCR data by the comparative CT method. Nat Protoc 3:1101–1108.

39. Planck KA, Rhee K. 2021. Metabolomics of Mycobacterium tuberculosis, p. 579–593. In Parish, T, Kumar, A (eds.), Methods in Molecular Biology, Springer Protocols. Springer Nature.

40. Banerjee D, Menasalvas J, Chen Y, Gin JW, Baidoo EEK, Petzold CJ, Eng T, Mukhopadhyay A. 2025. Addressing genome scale design tradeoffs in Pseudomonas putida for bioconversion of an aromatic carbon source. npj Syst Biol Appl 11:8.

41. Atkinson DE, Walton GM. 1967. Adenosine Triphosphate Conservation in Metabolic Regulation RAT LIVER CITRATE CLEAVAGE ENZYME. J Biol Chem 242:3239–3241.

42. Takaki K, Davis JM, Winglee K, Ramakrishnan L. 2013. Evaluation of the pathogenesis and treatment of Mycobacterium marinum infection in zebrafish. Nat Protoc 8:1114–1124.

43. Wayne L, Hayes L. 1996. An In Vitro Model for Sequential Study of Shiftdown of Mycobacterium tuberculosis through Two Stages of Nonreplicating Persistence. INFECTION AND IMMUNITY 64:2062–2069.

44. Lien KA, Dinshaw K, Nichols RJ, Cassidy-Amstutz C, Knight M, Singh R, Eltis LD, Savage DF, Stanley SA. 2021. A nanocompartment system contributes to defense against oxidative stress in Mycobacterium tuberculosis. eLife 10:e74358.

45. Gauthier DT, Doss JH, LaGatta M, Gupta T, Karls RK, Quinn FD. 2022. Genomic Degeneration and Reduction in the Fish Pathogen Mycobacterium shottsii. Microbiol Spectr 10:e01158–21.

46. Das S, Pettersson BMF, Behra PRK, Mallick A, Cheramie M, Ramesh M, Shirreff L, DuCote T, Dasgupta S, Ennis DG, Kirsebom LeifA. 2018. Extensive genomic diversity among Mycobacterium marinum strains revealed by whole genome sequencing. Sci Rep 8:12040.

47. Doig KD, Holt KE, Fyfe JA, Lavender CJ, Eddyani M, Portaels F, Yeboah-Manu D, Pluschke G, Seemann T, Stinear TP. 2012. On the origin of Mycobacterium ulcerans, the causative agent of Buruli ulcer. BMC Genom 13:258.

48. Stinear TP, Seemann T, Pidot S, Frigui W, Reysset G, Garnier T, Meurice G, Simon D, Bouchier C, Ma L, Tichit M, Porter JL, Ryan J, Johnson PDR, Davies JK, Jenkin GA, Small PLC, Jones LM, Tekaia F, Laval F, Daffé M, Parkhill J, Cole ST. 2007. Reductive evolution and niche adaptation inferred from the genome of Mycobacterium ulcerans, the causative agent of Buruli ulcer. Genome Res 17:192–200.

49. Brown AC, Parish T. 2008. Dxr is essential in Mycobacterium tuberculosisand fosmidomycin resistance is due to a lack of uptake. BMC Microbiol 8:78.

50. Gill WP, Harik NS, Whiddon MR, Liao RP, Mittler JE, Sherman DR. 2009. A replication clock for Mycobacterium tuberculosis. Nat Med 15:211–214.

51. Brown A, Kokoczka R, Parish T. 2015. LytB1 and LytB2 of Mycobacterium tuberculosis Are Not Genetically Redundant. Plos One 10:e0135638.

52. Goldstein JL, Brown MS. 1990. Regulation of the mevalonate pathway. Nature 343:425–430.

53. Salari S, Lee H, Tsantrizos YS, Park J. 2024. Inhibition of human mevalonate kinase by allosteric inhibitors of farnesyl pyrophosphate synthase. FEBS Open Bio 14:1320–1339.

54. Banerjee A, Wu Y, Banerjee R, Li Y, Yan H, Sharkey TD. 2013. Feedback Inhibition of Deoxy-d-xylulose-5-phosphate Synthase Regulates the Methylerythritol 4-Phosphate Pathway*. J Biol Chem 288:16926–16936.

55. Dorsey JK, Porter JW. 1968. The Inhibition of Mevalonic Kinase by Geranyl and Farnesyl Pyrophosphates. J Biol Chem 243:4667–4670.

56. Hinson DD, Chambliss KL, Toth MJ, Tanaka RD, Gibson KM. 1997. Post-translational regulation of mevalonate kinase by intermediates of the cholesterol and nonsterol isoprene biosynthetic pathways. J Lipid Res 38:2216–2223.

57. Voynova NE, Rios SE, Miziorko HM. 2004. Staphylococcus aureus Mevalonate Kinase: Isolation and Characterization of an Enzyme of the Isoprenoid Biosynthetic Pathway. J Bacteriol 186:61–67.

58. Lee YN, Colston MJ. 1986. The measurement of adenylate energy charge (AEC) in mycobacteria, including Mycobacterium leprae. FEMS Microbiol Lett 35:279–281.

59. Thomson M, Liu Y, Nunta K, Cheyne A, Fernandes N, Williams R, Garza-Garcia A, Larrouy-Maumus G. 2022. Expression of a novel mycobacterial phosphodiesterase successfully lowers cAMP levels resulting in reduced tolerance to cell wall–targeting antimicrobials. J Biol Chem 298:102151.

60. Wiese M, Seydel U. 1995. Monitoring of drug effects on cultivable mycobacteria and Mycobacterium leprae via the determination of their adenylate energy charges (AEC). J Microbiol Methods 24:65–80.

61. Madigan CA, Cambier CJ, Kelly-Scumpia KM, Scumpia PO, Cheng T-Y, Zailaa J, Bloom BR, Moody DB, Smale ST, Sagasti A, Modlin RL, Ramakrishnan L. 2017. A Macrophage Response to Mycobacterium leprae Phenolic Glycolipid Initiates Nerve Damage in Leprosy. Cell 170:973–985.e10.

62. Huang S, Zhou W, Tang W, Zhang Y, Hu Y, Chen S. 2021. Genome-scale analyses of transcriptional start sites in Mycobacterium marinum under normoxic and hypoxic conditions. BMC Genom 22:235.

63. Lin C, Tang Y, Wang Y, Zhang J, Li Y, Xu S, Xia B, Zhai Q, Li Y, Zhang L, Liu J. 2022. WhiB4 Is Required for the Reactivation of Persistent Infection of Mycobacterium marinum in Zebrafish. Microbiol Spectr 10:e00443–21.

64. Batra PP, Rilling HC. 1964. On the mechanism of photoinduced carotenoid synthesis: Aspects of the photoinductive reaction. Arch Biochem Biophys 107:485–492.

65. Mathews MM. 1963. STUDIES ON THE LOCALIZATION, FUNCTION, AND FORMATION OF THE CAROTENOID PIGMENTS OF A STRAIN OF MYCOBACTBRIUM MARINUM. Photochem Photobiol 2:1–8.

66. Ramakrishnan L, Tran HT, Federspiel NA, Falkow S. 1997. A crtB homolog essential for photochromogenicity in Mycobacterium marinum: isolation, characterization, and gene disruption via homologous recombination. J Bacteriol 179:5862–5868.

67. Mann FM, Xu M, Davenport EK, Peters RJ. 2012. Functional characterization and evolution of the isotuberculosinol operon in Mycobacterium tuberculosis and related Mycobacteria. Front Microbiol 3:368.

68. Bitok JK, Meyers CF. 2012. 2C-Methyl-d-erythritol 4-Phosphate Enhances and Sustains Cyclodiphosphate Synthase IspF Activity. ACS Chem Biol 7:1702–1710.

69. Provvedi R, Kocíncová D, Donà V, Euphrasie D, Daffé M, Etienne G, Manganelli R, Reyrat J-M. 2008. SigF Controls Carotenoid Pigment Production and Affects Transformation Efficiency and Hydrogen Peroxide Sensitivity in Mycobacterium smegmatis. J Bacteriol 190:7859–7863.

70. Hümpel A, Gebhard S, Cook GM, Berney M. 2010. The SigF Regulon in Mycobacterium smegmatis Reveals Roles in Adaptation to Stationary Phase, Heat, and Oxidative Stress. J Bacteriol 192:2491–2502.

71. Hartkoorn RC, Sala C, Uplekar S, Busso P, Rougemont J, Cole ST. 2012. Genome-Wide Definition of the SigF Regulon in Mycobacterium tuberculosis. J Bacteriol 194:2001–2009.

72. Yang C, Gao X, Jiang Y, Sun B, Gao F, Yang S. 2016. Synergy between methylerythritol phosphate pathway and mevalonate pathway for isoprene production in Escherichia coli. Metab Eng 37:79–91.

